# An RNA-centric atlas of human cell cycle regulation

**DOI:** 10.64898/2026.11.01.763428

**Authors:** Giulia Guiducci, Alice O Coomer, Dmitry Prokopov, Eleni Maniati, Charlie Brett, Parnia Babaei, María del Carmen Rodríguez Rodríguez, Martin Dodel, Anisha Thind, Jun Wang, Faraz K Mardakheh, Ozgen Deniz, Lovorka Stojic

## Abstract

Cell cycle progression is orchestrated by protein networks coordinating cell growth and division, yet the role of RNA-mediated mechanisms remains underexplored. Here, we define an RNA-centric landscape of human cell cycle control using palbociclib-synchronised normal diploid cells, profiling long non-coding RNAs (lncRNAs), transposable elements (TEs), alternative splicing (AS), and RNA-binding proteins (RBPs). We identify 104 cell cycle-regulated lncRNAs and validate lncRNA *ENSG00000230615,* whose perturbation alters cell cycle progression and downregulates neighbouring *KLF17*. Across the transcriptome, 297 TE loci exhibit differential expression dynamics during the cell cycle, with Alu elements upregulated during the G1-to-S transition. Cell-cycle-dependent AS changes affect 598 genes, with alternative 5’ splice sites showing the largest shift in usage at the G1-to-S transition. Furthermore, we show that cytokinesis proteins act as non-canonical RBPs with RNA-dependent stability. This study provides a comprehensive resource on RNA-based cell cycle control and highlights an underappreciated layer of regulation mediated by RNA mechanisms.

**Highlights:**

- First atlas of lncRNAs, transposable elements, alternative splicing and RNA-binding proteins across human cell cycle progression
- Only 104 lncRNAs show cell cycle regulation, independent of palbociclib treatment
- AluS expression and alternative 5’ splice site are enriched during the G1-to-S transition
- The cytokinesis protein PRC1 functions as a non-canonical RBP in interphase

## Introduction

Cell cycle control is a tightly regulated process and depends on two key events: accurate replication of genomic DNA and its faithful segregation into daughter cells, both of which occur in distinct phases of the cycle^1^. Error-free progression through the G1, S, G2, and M phases of the cell cycle requires tight regulation of gene transcription, translation, protein-protein interactions, post-translational modifications (PTMs), and ubiquitin-dependent protein degradation. These mechanisms collectively safeguard genome stability, and alterations in these pathways are closely linked to human disease, including cancer^2^. Despite decades of extensive research, RNA-mediated regulation underlying cell cycle control remains comparatively underexplored. Elucidating RNA-based mechanisms is therefore essential to fully understand cell cycle regulation and its dysregulation in disease.

Examples of RNA-mediated mechanisms include widely expressed long non-coding RNAs (lncRNAs), which mediate different stages of the cell cycle, either by regulating the expression of key cell cycle genes or by modulating the activities and stability of cell cycle proteins^3,4^. Alternative splicing (AS) produces different mRNA isoforms and, therefore, increases protein diversity. It can control the cell cycle by modifying the splicing of crucial cell cycle genes, or it can be regulated through PTMs of splicing factors^5,6^. Alongside AS and lncRNAs, transposable elements (TE)^7^, repetitive mobile genetic sequences that make up half of the human genome, represent an underexplored layer of cell cycle control. On the other hand, RNA-binding proteins (RBPs) control essentially all stages of RNA life, including transcription, splicing, RNA modifications, intracellular trafficking, translation, and RNA decay^8^. However, the catalogue of RBPs has expanded to include numerous non-canonical RBPs that lack classical RNA-binding domains and participate in riboregulation, a process in which RNA directly regulates protein activity ^9^. Riboregulation has recently been shown to play a key role in mitosis, with hundreds of mitotic proteins displaying RNA-dependent interactions, thereby adding an additional regulatory layer to cell cycle control^10^.

These RNA-based elements frequently co-operate; for example, many lncRNAs exert their functions by engaging multiple RBPs, leading to diverse molecular and cellular outcomes ^11^. While studies on the role of lncRNAs^12–15^, enhancer RNAs^16^, RBPs^17^, and, to a lesser extent, on AS^18^ and TE^19^ in cell cycle regulation have been performed, they have mostly relied on cancer cell lines, low numbers of biological replicates, and synchronisation strategies such as double thymidine block^12,14,15,20^, which leads to excessive DNA damage^21,22^ and chromosomal aberrations^23^. These limitations have restricted our understanding of RNA-based mechanisms in normal cell cycle physiology. To overcome these challenges, we used palbociclib, a CDK4/6 inhibitor that arrests cells in the G1 phase^24^, with minimal DNA damage and preservation of genome integrity^22,25^. In this study, we profiled the landscape of lncRNAs, TEs, AS and RBPs by performing transient palbociclib-induced synchronisation in a non-transformed near-diploid telomerase-immortalised hTERT-RPE1 cell line. We found novel cycle-regulated lncRNAs whose expression was independent of palbociclib treatment. Using a new TEgment pipeline (*Prokopov et al., 2026; manuscript co-submitted back-to-back*), we identified TE loci exhibiting cell cycle stage-specific expression, with standalone AluS elements significantly enriched at the G1-to-S phase transition. We also identified cell cycle-regulated AS events, with alternative 5’ splice site usage being significantly enriched at the G1-to-S phase transition. Lastly, by profiling RBP dynamics across the cell cycle using Orthogonal Organic Phase Separation (OOPS)^26^, we found that non-canonical RBPs, including cytokinesis proteins such as Protein Regulator of Cytokinesis 1 (PRC1), show RNA-dependent stability in interphase. Our work presents the first comprehensive RNA-centric atlas of human cell cycle regulation, providing a valuable resource for uncovering previously underappreciated regulatory mechanisms.

## Results

### Efficient synchronisation of RPE1 cells with transient palbociclib treatment enables profiling of RNA landscape during cell cycle progression

To systematically investigate the roles of lncRNAs, TEs, AS, and RBPs during cell cycle progression, we used telomerase-immortalised hTERT-RPE1 cells (hereafter, RPE1), a diploid, non-transformed retinal epithelial cell line with intact cell cycle control and widely used in cell cycle studies^22,25^. RPE1 cells were synchronised by transient exposure to 500 nM palbociclib^22^, which arrests cells at the G1 phase, preventing cell cycle progression. Following drug washout, samples for fluorescence-activated cell sorting (FACS), RNA, and protein analysis were collected every 2-3 hours (Figure 1A). FACS analysis confirmed that palbociclib induces G1 arrest and enables highly synchronous progression after drug release, as shown by cell cycle profiles across time points (T): T0 (G1 arrest), T2-T5 (G1/S transition), T5–T8 (mid/late S phase), and finally into T11-T12 (G2/M phase) (Figures 1B, 1C). The elevated mRNA and protein levels of *Cyclin E1* and *Cyclin B1*, key markers for the G1/S and G2/M cell cycle stages, respectively, support the timing of these cell cycle transitions (Figures 1D, 1E). This confirms successful synchronisation of RPE1 cells using palbociclib.

**Figure 1.**
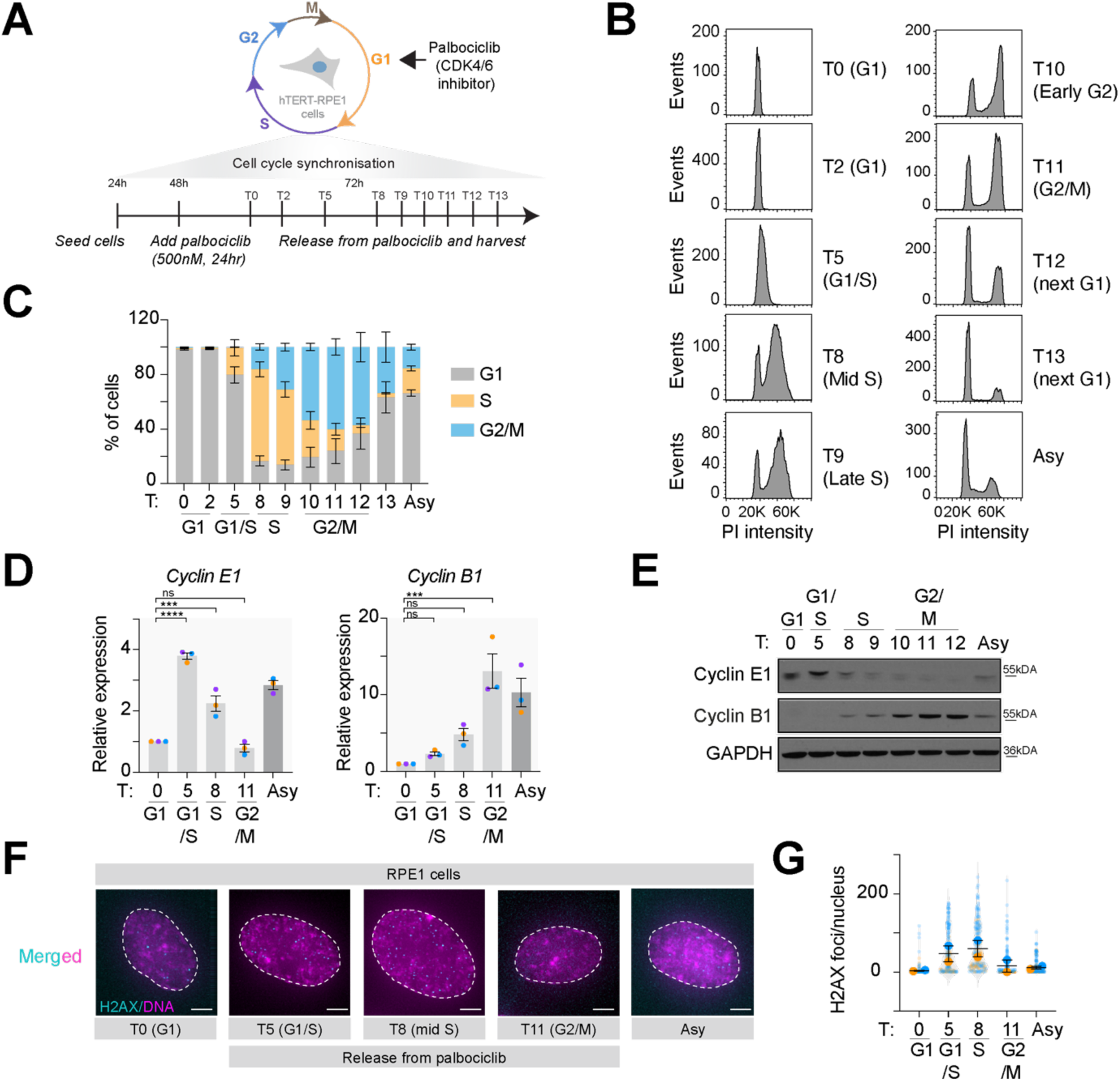
Transient palbociclib treatment enables highly synchronous cell cycle progression in RPE1 cells. (A) Scheme showing experimental design of RPE1 cells synchronisation using the palbociclib, the CDK4/6 inhibitor and timed releases with sample collection. (B) Representative RPE1 FACS profiles with corresponding quantifications (C) at indicated time points (expressed in hours) after release from 500 nM palbociclib treatment. N = 5. (C) Quantification of relative cell cycle stages as in panel (B) for the indicated time points following the Palbociclib release in RPE1 cells. N = 5. (D) Relative expression of *Cyclin E1* (left) and *Cyclin B1* (right) at selected time points following palbociclib release in RPE1 cells, as measured by qPCR. Results are presented relative to unreleased cells (T0, G1 arrest). N = 3. (E) Western blot showing the levels of Cyclin E1 and Cyclin B1 at the indicated time points (hours) following Palbociclib release in RPE1 cells. GAPDH was used as a loading control. (F) Representative images of interphase RPE1 cells stained for the DNA damage marker γH2AX (cyan) and DNA (Hoescht, magenta) at indicated time points (hours) following release from palbociclib treatment. Scale bar, 5 μm. (G) Quantification of γH2AX staining was determined by quantifying the number of γH2AX foci per nucleus (F). N = 2. (n_(T0)=_160, n_(T5)=_162, n_(T8)=_161, n_(T11)=_162, n_(Asy)=_160). Error bars in all panels are shown as mean ± S.E.M. Scale bar, 5 μm. One-way ANOVA with Dunnett’s multiple comparison test was used in panel (D) to determine statistical significance: ns, not significant; ***p < 0.001; ****p < 0.0001. Asy denotes asynchronous cells.

To avoid DNA damage associated with common synchronisation methods, such as the double thymidine block^21,23^, we quantified the frequency of foci containing phosphorylated γH2AX at serine 139 (a DNA damage marker) to assess DNA damage after palbociclib treatment and after release from arrest. Consistent with previous reports^22,25^, γH2AX foci appeared during S phase (T5, T8), reflecting physiological replication-associated DNA damage^27,28^, while G1-enriched (T0) and G2/M-enriched (T11) cells showed minimal or no γH2AX signal (Figures 1F, 1G). Together, these results demonstrate that transient palbociclib synchronisation yields synchronous RPE1 populations without inducing substantial DNA damage, providing a suitable model for profiling the RNA landscape across the cell cycle.

### Transcriptome analysis of RPE1 cells at different stages of the cell cycle

We leveraged the established RPE1 synchronisation to profile transcriptomic changes across cell cycle stages by performing bulk RNA sequencing (RNA-seq) at representative time points (T0, T5, T8 and T11) in four biological replicates. This dataset, which combines synchronous, non-transformed cells with robust reproducibility, provides greater statistical power and a more physiologically relevant framework for studying cell cycle-dependent RNA dynamics than many existing public datasets^12–14,16^ (Figure S1A). To identify cell cycle stage-specific genes, we defined differentially expressed genes (DEGs) as those that (i) showed the highest transcripts per million (TPM) expression in one phase relative to the other phases and (ii) were significantly differentially expressed in the current phase relative to the adjacent phase (Benjamini–Hochberg, padj < 0.05 and |log2FC| > 0.5). Using these criteria, we identified 1,116 DEGs across the cell cycle (Figure 2A, Table S1). Notably, protein-coding genes upregulated in the G1 and G1/S stages of the cell cycle were enriched for signal transduction, transcription and response to stimuli; those upregulated in the S phase were enriched for DNA replication and chromatin remodelling; and those upregulated in the G2/M phase were enriched for mitosis and microtubule organisation (Figure S1B-S1E, Table S2), consistent with well-established stage programmes. Of the 1,116 DEGs, only 104 were annotated as lncRNA genes in our dataset (Figure 2B, Table S1), approximately ten times fewer than previously reported^12^ (Figure S1F). This discrepancy is likely attributable to differences in synchronisation strategy and cellular models (Figure S1A).

**Figure 2.**
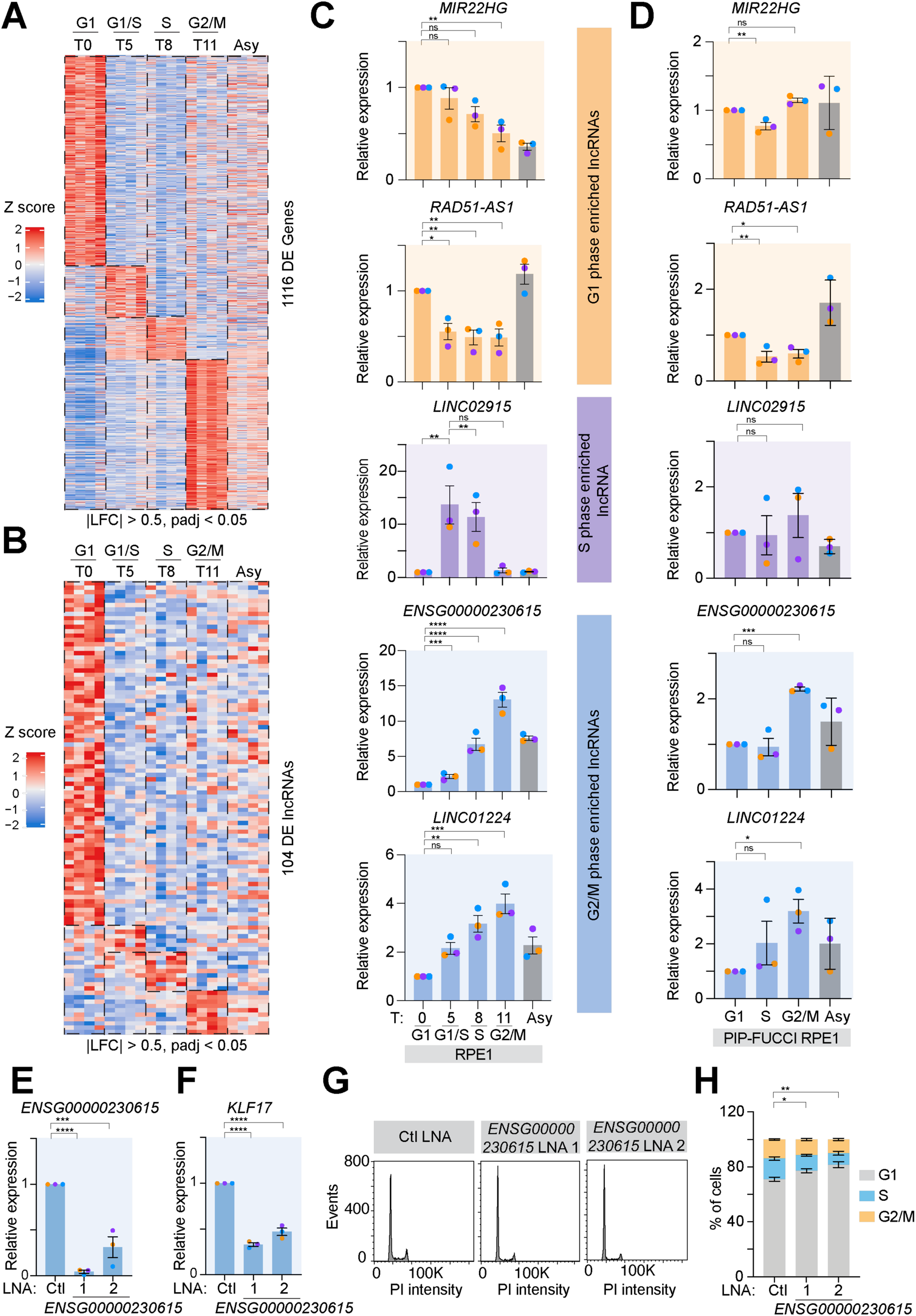
Identification of cell cycle-regulated lncRNAs reveals an uncharacterised lncRNA *ENSG00000230615* as a regulator of cell cycle progression. (A) Heatmap of normalised log_2_TPM expression of all cell cycle phase-specific genes identified by RNA-sequencing (highest expression in cycle phase & |log_2_ fold change| ≥ 0.5 with BH-adjusted *p* ≤ 0.05 compared to the adjacent cell-cycle phase). Time points (hours) following release from palbociclib treatment are indicated. N = 4. (B) Heatmap of normalised log_2_TPM expression of cell cycle phase-specific lncRNAs, extracted from the dataset shown in (A). (C) Validation of the top five cell cycle-regulated lncRNAs in RPE1 cells. Relative expression of G1-phase (*RAD51-AS1*, *MIR22HG*; orange), S-phase (*LINC02915*; purple), and G2/M-phase regulated (*ENSG00000230615*, *LINC01224*; blue) lncRNAs at indicated time points (hours) following palbociclib release, as measured by qPCR. Results are presented relative to unreleased cells (T0, G1 arrest). N = 3. (D) Validation of the top five cell cycle-regulated lncRNAs in PIP-FUCCI RPE1 cells. Relative expression of G1-phase (*RAD51-AS1*, *MIR22HG*; orange), S-phase (*LINC02915*; purple), and G2/M-phase regulated (*ENSG00000230615*, *LINC01224*; blue) lncRNAs at indicated cell cycle stages, as measured by qPCR. Results are presented relative to G1 samples. N = 3. (E, F) *ENSG00000230615* (E) and *KLF17* (F) expression in RPE1 cells following depletion with two LNA Gapmers targeting exon 2 (LNA 1) and exon 3 (LNA 2), as measured by qPCR. Results are presented relative to the negative control LNA (LNA Ctl). N = 3. (G, H) Representative RPE1 cell cycle profiles (G) with corresponding quantifications (H) following depletion of *ENSG00000230615* lncRNA using the LNA Gapmers described in (E). N = 5. Error bars in (C), (D), (E), (F) and (H) are shown as mean ± S.E.M.One-way ANOVA with Dunnett’s multiple comparisons test was used in panels (C), (E) and (G) to determine statistical significance: ns, not significant; *p < 0.05; **p < 0.01; ***p < 0.001; ****p < 0.0001. Asy denotes asynchronous cells.

Next, we prioritised stage-specific lncRNA candidates for validation using four criteria: (i) conservation at the synteny level (Figures S2A-S2D), (ii) proximity to relevant protein-coding genes with known cell cycle roles such as RAD51^29^ and KLF17^30^ (Figures S2B-S2D), (iii) cancer relevance based on dysregulation (Figures S3A-S3E, Table S3) and correlation with patients’ survival (Figures S4A-S4G) and (iv) limited prior characterisation. Applying these criteria, we selected *MIR22HG* and *RAD51-AS1* (G1-enriched), *LINC02915* (S-enriched), *ENSG00000230615* and *LINC01224* (both G2/M-enriched). Genomic loci for each candidate are shown in Figures S2A–S2E. Coding potential analysis by CPC2^31^ indicated noncoding status for all candidates except *LINC02915*, which exhibited significant coding potential (Figure S3F). Cell cycle regulation of selected lncRNAs was confirmed in an independent set of biological replicates in RPE1 cells (Figure 2C). To exclude palbociclib-specific effects, we validated their phase-specific expression using RPE1 cells expressing the PIP-FUCCI (Fluorescent Ubiquitination-based Cell Cycle Indicator) reporter, which allows better cell cycle phase reporting based on the PIP degron (PCNA-interacting protein) from the human Cdt1 (Figures S5A, S5B)^32^. In PIP-FUCCI RPE1 cells, *MIR22HG, RAD51-AS1, ENSG00000230615*, and *LINC01224* showed the expected phase-specific expression, while *LINC02915* did not validate (Figure 2D). This likely reflects higher *LINC02915* expression at the G1-to-S transition (T5) in RPE1 cells (Figure 2C), a phase that is not well captured by the PIP-FUCCI RPE1 system^32^.

Because *ENSG00000230615* was validated in both RPE1 models, it is a poorly characterised syntenically conserved lncRNA and lies adjacent to KLF17, a transcription factor with tumour-suppressive activity and reported effects on cell cycle progression in cancer^30,33^, we prioritised it for functional characterisation. Subcellular fractionation revealed that *ENSG00000230615* is a chromatin-enriched lncRNA in RPE1 cells (Figure S5C), consistent with ENCODE cell line data (Figure S5D). Given its nuclear localisation, we employed Locked Nucleic Acid (LNA) Gapmers, which are highly efficient at depleting nuclear lncRNAs^34^. Depletion of *ENSG00000230615* with two independent LNA gapmers targeting different exons led to downregulation of *KLF17* (Figure 2E, 2F) and G1 cell cycle arrest (Figure 2G, 2H). These data suggest that lncRNA *ENSG00000230615* may regulate the cell cycle by modulating *KLF17* expression in *cis*.

### TEgment reveals cell cycle stage-specific TE distributions and enrichment of standalone AluS elements at the G1-to-S transition

We sought to assess the cell-cycle regulatory potential of TEs by leveraging our RNA-seq dataset with a newly established pipeline, TEgment (*Prokopov et al., 2026; manuscript co-submitted back-to-back*). This pipeline allows for locus-specific identification, quantification, and structural classification of TEs from transcriptome assemblies while minimising intronic noise. We defined cell cycle-regulated TE loci as those differentially expressed between adjacent cell cycle phases (|log_2_FC| > 0.5 and adjusted p < 0.05), yielding 297 cell cycle-regulated TE loci (Figure 3A, Table S4). Unsupervised hierarchical clustering divided these loci into four distinct TE clusters: cluster 1, TE loci with the highest expression in G1 phase; cluster 2, TE loci with the highest expression at the G1-to-S transition; cluster 3, TE loci with the highest expression in mid-to-late S phase; cluster 4, TE loci with the highest expression in the G2/M phase (Figure 3B). Classifying the loci revealed distinct distributions of TEs across clusters. For example, lncRNA-embedded TEs were enriched in G1 phase (cluster 1), standalone TEs were enriched at G1-to-S transition (cluster 2), while unclassified TEs predominated in mid–late S and G2/M phase of the cell cycle (clusters 3–4).

**Figure 3.**
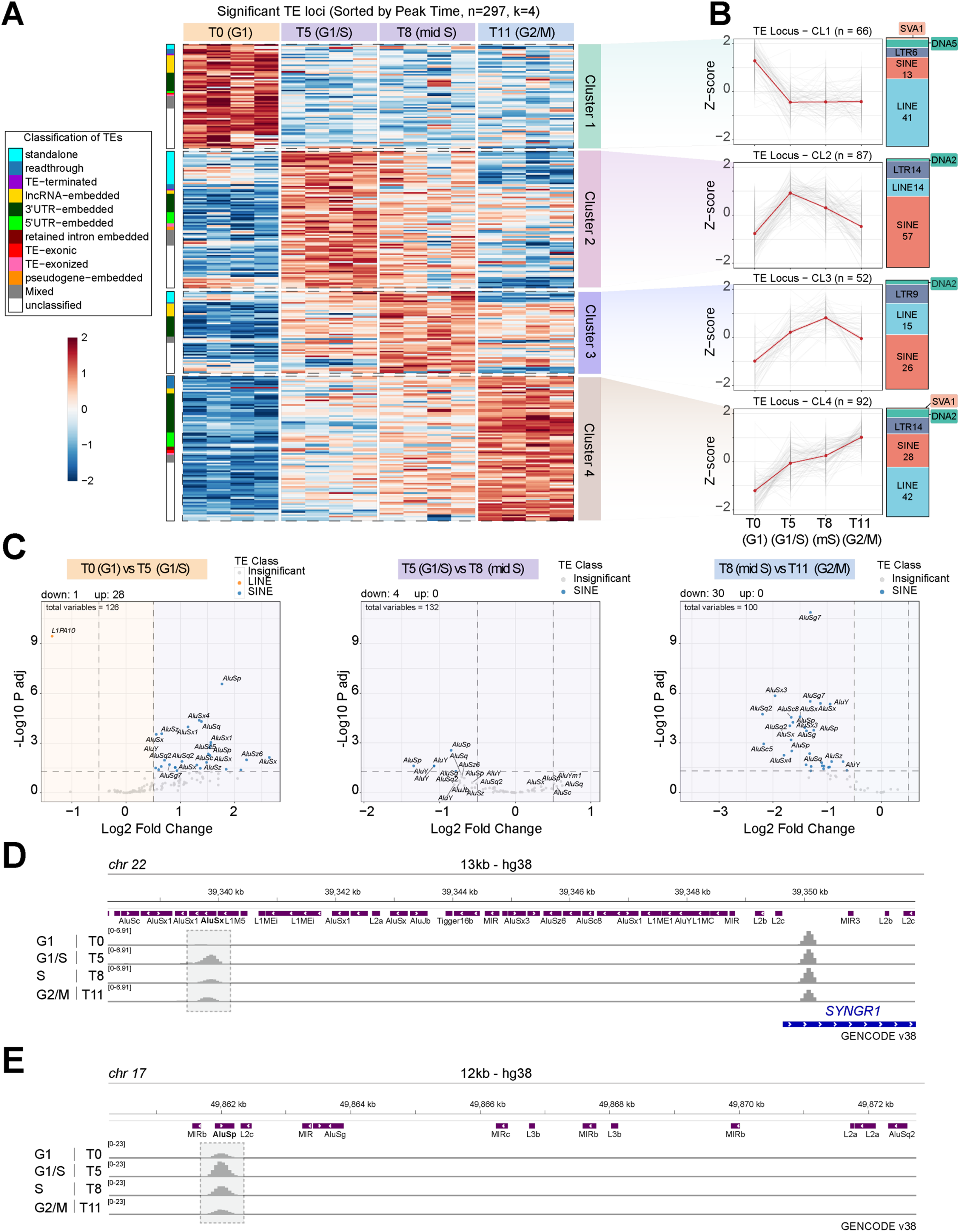
Expression profiling and classification of cell cycle-regulated transposable elements reveal an enrichment of standalone AluS elements at the G1-to-S transition. (A) Heatmap (k = 4) of VST expression values of cell cycle phase-specific TEs identified by TEgment (|fold change| ≥ 0.5 and adjusted *p* ≤ 0.05, compared with the adjacent cell-cycle phase). Time points (hours) following release from palbociclib treatment are indicated. Asy denotes asynchronous cells. Classification of TE elements, performed with TEgment, is displayed on the left. N = 4. (B) Trend plots of z-normalised VST values for TE loci in each cluster defined in (A), red lines represent median z-values. Bar plots on the right show the composition of each cluster by TE superfamilies (DNA in green; LINE in cyan; LTR in; SINE in red; SVA in salmon). mS=mid S phase. (C) Volcano plots of differentially expressed standalone TEs across adjacent phases (dashed lines indicate |fold change| ≥ 0.5 and adjusted *p* ≤ 0.05). Both significant TE superfamilies and individual loci are shown (LINE in orange; SINE in blue; non-significant in grey). (D) Representative example of G1/S-specific upregulation of an AluS element located in proximity to a protein-coding gene. Snapshot was taken from the IGV genome browser, including GENCODE v38 and RepeatMasker annotations. (E) Representative example of G1/S-specific upregulation of an AluS element located in an intergenic region, with no nearby protein-coding genes. Snapshot was taken from the IGV genome browser, including GENCODE v38 and RepeatMasker annotations.

We then examined the TE class composition across clusters and found distinct profiles in each cell cycle stage. LINE elements predominated in G1 and G2/M (cluster 1 and 4, respectively), whereas SINE elements dominated the G1/S and S phase (cluster 2 and 3, respectively) (Figure 3B). LTR and DNA transposons represented only minor fractions across all clusters. The strong overrepresentation of SINEs and standalone TEs, elements activated by autonomous promoters, prompted us to focus on G1/S–associated TE loci (cluster 2). We found intriguing occurrences in which all cell cycle-regulated TEs within the standalone category belonged to the AluS subfamily, a primate-specific class of intermediate-age Alu elements^35,36^. These AluS loci are upregulated at the G1-to-S transition, remain expressed across S phase and are downregulated in the G2/M stage of the cell cycle (Figure 3C). Representative examples (Figure 3D, 3E) show that AluS can be upregulated independently of nearby protein-coding genes or other repeats, consistent with autonomous regulation. Although systematic profiling of TE expression throughout the cell cycle has been lacking, prior overexpression studies suggested that AluS elements can influence S-phase progression in a non-tumour context^19^. Our findings corroborate this initial evidence and additionally reveal stage-specific distributions of cell cycle-regulated TEs across categories and superfamily.

### Alternative splicing is dynamically regulated across the cell cycle with increased alternative 5’ splice site usage at the G1-to-S transition

To systematically investigate alternative splicing across the cycle, we applied the rMATS^37^ pipeline to analyse differential splicing on our RNA-seq dataset. Cell-cycle-regulated splicing events were defined as those with the highest average percentage splicing inclusion (PSI) in one phase and significant differences relative to the adjacent cell cycle phase (FDR < 0.05 and |dPSI| > 0.1). Overall, we identified 717 cell cycle-regulated splicing events mapping to 654 differentially spliced genes (Figure 4A, Table S5). Of 654 differentially spliced genes, only 56 overlapped with cell cycle-regulated differentially expressed genes (Figure 4B), indicating that alternative splicing is largely uncoupled from changes in transcript abundance and represents an additional post-transcriptional layer of cell cycle regulation, in agreement with previous work^18^. Gene Ontology (GO) analysis of the 598 differentially spliced genes revealed strong enrichment for pathways involved in nucleic acid processes, microtubule cytoskeletal organisation, and DNA repair (Figure 4C, Table S6).

**Figure 4.**
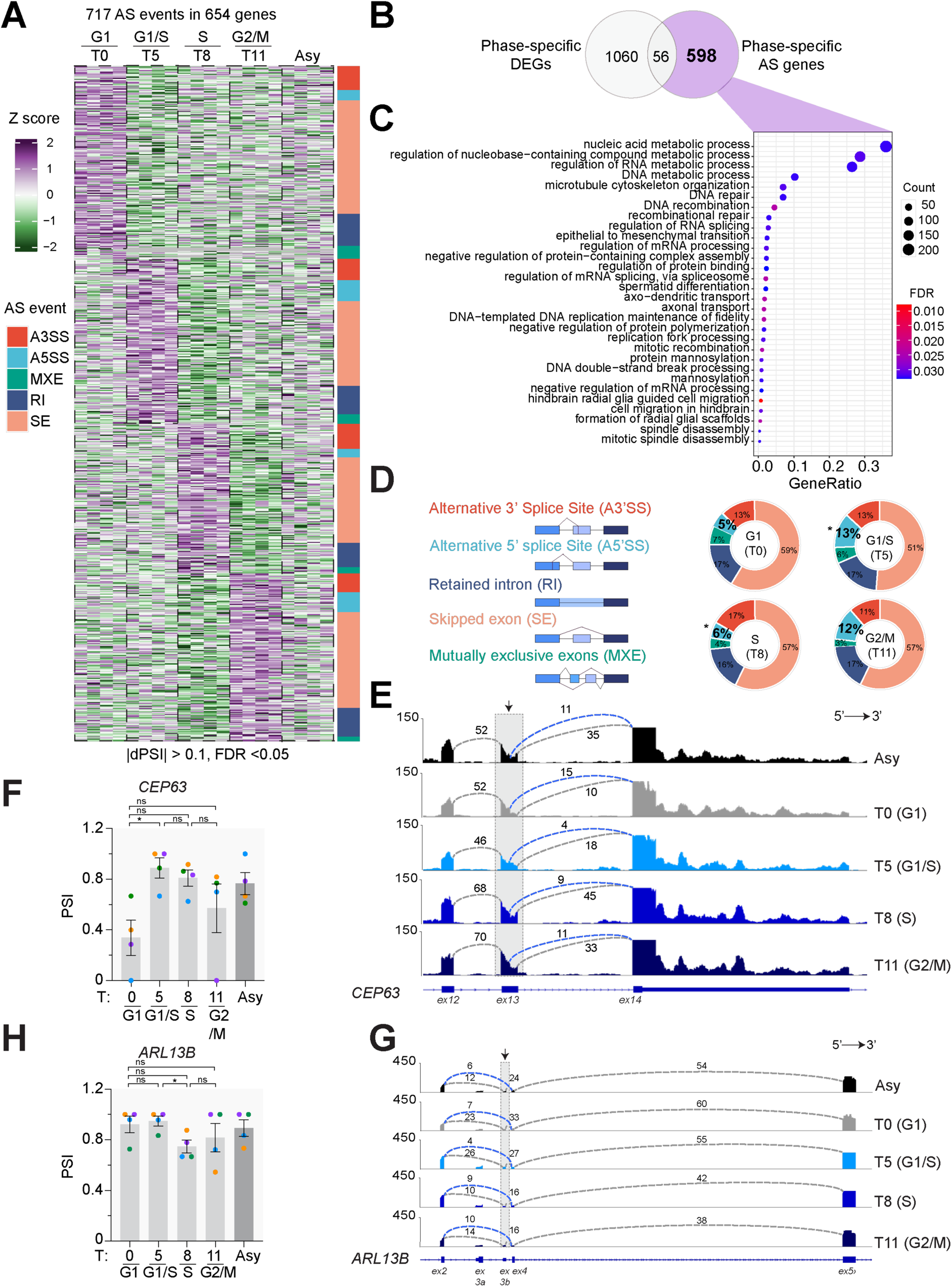
Cell cycle-regulated alternative splicing events show increased alternative 5’ splice site usage at the G1-to-S transition. (A) Heatmap of inclusion levels for cycle phase-specific alternatively splicing events identified by rMATS (splicing events with the highest average inclusion level in each cycle phase & |dPSI| ≥ 0.1 with FDR ≤ 0.05 compared to the adjacent cell-cycle phase). Time points (hours) following palbociclib are indicated. The classification of splicing events is shown on the right. A3’SS: alternative 3’ splice sites, A5’SS: alternative 5’ splice sites, SE: skipped exon, MXE: mutually exclusive exons, RI: retained intron. N = 4. (B) Venn diagrams intersecting DEGs across the cell cycle from Fig. 2A (grey) with genes undergoing alternatively splicing (purple). (C) Gene Ontology (GO) analysis of the 598 phase-specific alternatively spliced genes that are not differentially expressed. The top 30 most enriched terms are shown (D) Doughnut charts showing the percentage distribution of splicing events across cell cycle stages. *p < 0.05, chi-square test compared to adjacent cycle phase. (E) Sashimi plot of *CEP63* exon 13 alternative 5’ splice site (the relevant exon is highlighted with a grey rectangle) in time points (hours) following palbociclib release. (F) Percentage splicing inclusion (PSI) of *CEP63* exon 13 calculated from RNA-seq data (as shown in E). (G) Sashimi plot of *ARL13B* exon 3b skipping (the relevant exon is highlighted with a grey rectangle) in time points (hours) following palbociclib release. (H) PSI of *ARL13B* exon 3b, calculated from RNA-seq data at time points following palbociclib release (as shown in G). Error bars in (F) and (H) are shown as mean ± S.E.M. One-way ANOVA with Dunnett’s multiple comparisons test was used in panels (F), and (H) to determine statistical significance: ns, not significant; *p < 0.05. The tracks (E, G) were constructed from averages of four biological replicates for each condition (Asy in black, T0 in grey, T5 in cyan, T8 in blue, T11 in dark blue). Asy denotes asynchronous cells.

Analysis of cell cycle-dependent splicing events revealed that skipped exons (SE) are the most abundant events across all phases, followed by retained introns (RI), alternative 3’ splice sites (A3’SS), alternative 5’ splice sites (A5’SS) and mutually exclusive exons (MXE) (Figure 4D, Table S5). Notably, A5’SS was the only significant event that increased at the G1-to-S transition and then declined again from mid S phase (Figure 4D), suggesting a specific role in S-phase progression. Detailed evaluation of A5’SS events revealed that *CEP63*, which encodes a centrosomal protein required for centriole duplication^38,39^, undergoes an A5’SS at G1-to-S transition (T5), producing a longer exon (Figures 4E, 4F, and Figure S6A). This extended exon encodes a coiled-coil domain required for interaction with CEP152^40^ (Figure S6B). Given the essential role of the CEP63–CEP152 interaction in centriole duplication during S phase^38,39^, the alternative CEP63 isoform is likely biologically important for this process.

Among skipping exon events, *ARL13B*, a gene primarily known for roles in ciliogenesis^41^, undergoes mid-S (T8) exon skipping (Figures 4G, 4H, and Figure S6C). Inclusion of *ARL13B* exon 3b introduces a premature stop codon, producing a truncated isoform, whereas exon skipping in mid-S yields the full-length isoform. Although the precise role of ARL13B in cell cycle regulation remains unclear, previous research has linked ARL13B dysfunction to reduced S phase entry and an accumulation of cells in the G2/M stage of the cell cycle^42^. This suggests that mid-S exon skipping may facilitate progression into G2/M phase. Altogether, these results show that stage-specific alternative splicing, including increased alternative 5’ splice site usage at the G1-to-S transition, change protein domain composition and may directly impact cell-cycle progression.

### Integrative analysis of transcription (co)factors and splicing regulators reveals coordinated control of cell cycle-dependent transcriptomic changes

Our data identified profound stage-specific changes in gene and TE expression, as well as in alternative splicing, across the cell cycle. We next used RNA-seq data to dissect the interplay between transcriptional (co)factors and splicing regulators, which likely coordinate these regulatory programmes at distinct cell-cycle stages. First, we extracted phase-specific transcription (co)factors (i.e., TFs and TcoFs) by intersecting DEGs (Figure 2A) with AnimalTFDB 4.0, a curated resource for TFs and TcoFs across animal species^43^. This analysis identified 147 phase-specific TFs and TcoFs across the cell cycle, with the largest number detected at the G1 stage (T0, n=63), followed by G2/M (T11, n=59), G1/S (T5, n=20), and mid-S phase (T8, n=5) (Figure 5A-5D). The zinc finger C2H2 (zf-C2H2) family was the most abundant across phases, consistent with the large size of this family in humans ^44^ (Figure 5E). Second, we found that each phase was characterised by distinct enriched T(co)F families. For instance, the G1 phase contained the most exclusive families, including GTF2I, TSC22, SRF, Runt, and Forkhead (Figure 5A). The G1/S transition was enriched for AP-2, ESR-like, IRF5, and P53 family member p73 (Figure 5B), while the mid-S phase was marked by SMAD7, ZBTB14 and ID2, the latter playing a key role in cell cycle regulation by promoting entry into S phase and cell proliferation^45^ (Figure 5C). Furthermore, G2/M exhibited enrichment of the E2F5, PAX6, OTX2, THAP3, and zf-GATA families, indicating transcriptional reprogramming before mitosis (Figure 5D). Interestingly, OTX2 has been shown to be expressed during the G2/M phase of the cell cycle^46^, and E2F5, a component of the mammalian DREAM complex, is required to repress G1/S and G2/M cell cycle genes^47^, further supporting our findings.

**Figure 5.**
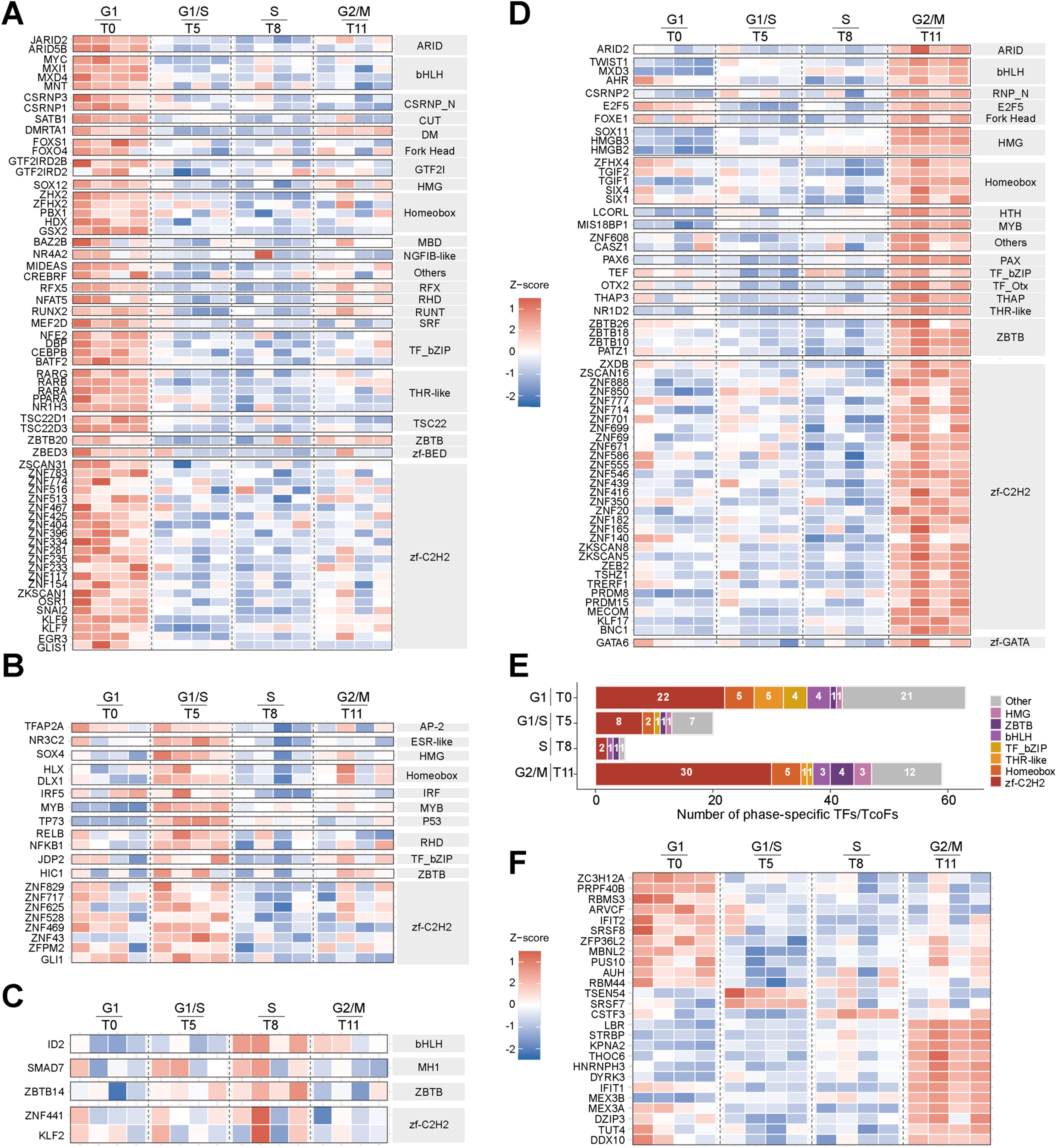
Integrated map of phase-specific regulators controlling gene expression, transposable elements and alternative splicing across the cell cycle. (A-D) Heatmap of z-score normalised expression (log_2_TPM) of phase-specific transcription (co)factors identified by RNA-seq (as in Figure 2A). Z-scores are calculated per gene across all samples, with replicates shown (N = 4). Time points (hours) following palbocilib release are indicated. Genes are ordered and grouped by TF family as annotated in AnimalTFDB^43^. Transcription (co)factors families represented include: ARID (AT-rich interaction domain), AP-2 (activating protein 2), bHLH (basic helix-loop-helix), CSRNP_N (cysteine-serine-rich nuclear protein), CUT (cut homeobox), DM (doublesex and mab-3 related), E2F (E2 factor), ESR-like (estrogen receptor-like), Fork_head (Forkhead), GTF2I (general transcription factor II-I), HMG (high mobility group), Homeobox, HTH (helix-turn-helix), IRF (interferon regulatory factor), MBD (methyl-CpG binding domain), MH1 (Mad homology 1), MYB (myeloblastosis), NGFIB-like (nerve growth factor IB-like), P53 (tumor protein p53), PAX (paired box), RFX (regulatory factor X), RHD (Rel homology domain), Runt (runt domain), SRF (serum response factor), TF_bZIP (basic leucine zipper), TF_Otx (orthodenticle homeobox), THAP (thanatos-associated protein), THR-like (thyroid hormone receptor-like), TSC22 (TSC22 domain family), ZBTB (zinc finger and BTB domain-containing), zf-BED (zinc finger BED domain), zf-C2H2 (zinc finger C2H2), and zf-GATA (zinc finger GATA domain). (E) Bar chart showing the number of transcription (co) factor genes from different families across the cell cycle (A-D). Each bar represents the total number of phase-specific differentially expressed T(co)F-encoding genes per phase, coloured by TF family. T(co)F families with four or fewer total counts across all phases are grouped as “Other”. (F) Heatmap of z-score normalised expression (log_2_TPM) of selected phase-specific splicing (co)regulators and RNA-binding proteins (RBPs) identified by RNA-seq (as in Figure 2A). Z-scores are calculated per gene across all samples, with replicates shown (N = 4). Time points (hours) following palbociclib release are indicated. Genes are grouped by peak phase and ranked by z-score within each phase. RBPs were identified by cross-referencing phase-specific DEGs against a curated reference list integrating GO term-based annotations (RNA splicing, splicing coactivator, and broader RNA-binding and processing categories) and experimentally validated RBPs from the RNA-Binding Protein Database^60^. The final gene selection was manually curated to retain only those with published evidence of a role in RNA splicing or in splicing regulation.

To capture transcriptional programs beyond phase-specific TF expression, we applied DoRothEA^48^ to infer TF activity from DEGs using curated TF-target interactions. This analysis confirmed expected regulators, including E2F1 at the G1/S transition^1^ and FOXM1 at G2/M^49^, and revealed activation of HIF1A and SMAD3 in the G1 phase, in line with previous studies ^50,51^(Figure S7A). We also observed E2F4 activation across S and G2/M phase, consistent with its emerging activator roles in addition to its canonical repressor function^52–54^ and FOXO3 activation before G1/S, concordant with its reported role in G1/S regulation^55^. Although we cannot fully exclude contributions from palbociclib treatment, our data indicate that distinct, phase-specific groups of transcription (co)factors shape the transcriptional landscape to drive orderly cell-cycle progression.

Given the strong enrichment of zf-C2H2 proteins, particularly Krüppel-associated box zinc finger proteins (KRAB-ZFP), which are established repressors of transposable elements^56^ and have been recently implicated in cell cycle regulation^57^, we asked whether specific KRAB-ZFPs might regulate the standalone AluS upregulation observed at S-phase onset (Figure 3C). Using KRABopedia, a catalogue of human KRAB-ZFP genomic targets^58^, we found that ZNF334, a G1-specific (T0) KRAB-ZFP (Figure 5A), is predominantly enriched at AluS elements (Figure S7B, S7C). By contrast, ZNF441, reported to target AluY elements^59^ and expressed in S phase (T8) (Figure 5C), was enriched at both AluY and AluS elements (Figure S7D, S7E). Although we cannot exclude contributions from other cell cycle-regulated KRAB-ZFPs^57^, we posit that the repressive activity of ZNF334 in G1 and of ZNF441 in S phase together might define the temporal windows for standalone AluS upregulation during S phase progression in RPE1 cells.

Lastly, we investigated potential drivers of A5’SS enrichment at G1-to-S transition (Figure 4D) by profiling RBPs involved in splicing regulation and/or RNA processing across cell-cycle stages. To address this, we intersected the DEGs with a manually curated list of splicing-related GO annotations (see Materials and Methods) and experimentally validated RBPs from the RNA-Binding Protein Database^60^, identifying 26 phase-specific regulators of RNA metabolism (Figure 5F). Only three displayed G1/S and S-phase-specific expression, with SRSF7 emerging as the prime candidate. Mechanistically, SRSF7 can modulate 5′ splice-site choice in a position-dependent manner. Indeed, when bound upstream of competing 5′ sites, it promotes U1 snRNP recruitment and favours the intron-proximal (downstream) site^61^, consistent with the increased A5′SS usage at G1-to-S transition. Notably, loss of SRSF7 has been reported to reduce S-phase entry in cancer cells^62^, and its G1/S phase enrichment in our data supports a potential functional link between SRSF7 activity and the observed A5′SS programme. Together, these findings suggest that phase-specific transcription factors and splicing regulators coordinate transcriptomic and splicing programmes across the cell cycle.

### OOPS identifies non-canonical RNA-binding proteins, including cytokinesis proteins, with dynamic RNA binding across the cell cycle

Having established dynamic regulation of lncRNAs, transposable elements and alternative splicing across the cell cycle, we next assessed the role of RBPs, a layer of post-transcriptional regulation examined to date in only isolated cases. To do this, we applied OOPS^26^, an unbiased method for purifying RNA-protein complexes from synchronised RPE1 cells (T0, G1; T5, G1/S; T8, mid-S; T11, G2/M) in parallel with total proteome profiling, followed by mass spectrometry. Normalising OOPS-derived RNA-bound peptide intensities to total proteome levels enabled the detection of changes in RNA-binding activity independent of protein abundance. We identified 45 proteins with altered RNA binding across cell cycle stages (Figure 6A, Table S7) and, strikingly, most are non-canonical RBPs that lack GO “RNA binding” annotations and canonical RNA-binding domains (Figure 6B), whose primary functional annotations relate to other cellular processes^9^.

**Figure 6.**
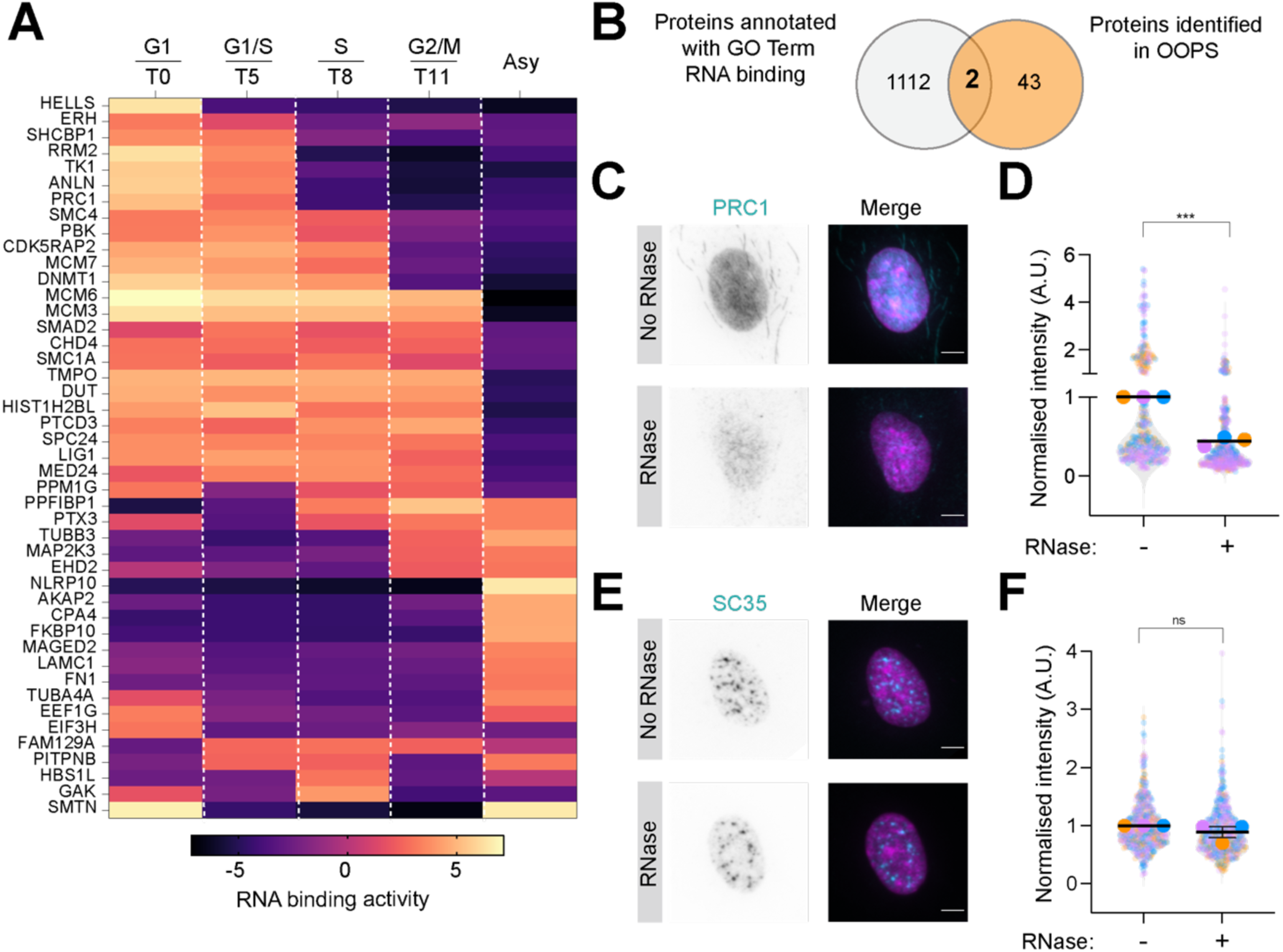
OOPS reveals dynamic, cell-cycle-dependent RNA binding by non-canonical RNA-binding proteins. (A) Heatmap of significant changes in the RNA-binding activity of RBPs in different phases of cell cycle. RNA-binding activity was calculated by normalising the OOPS-derived RNA-bound proteome intensities to whole cell lysate-derived total proteome intensities, from cells enriched in the indicated time points (hours) following release from palbociclib. Significance was calculated using a one-way ANOVA analysis coupled with post-hoc test (FDR < 0.05). The post-hoc scores were subjected to hierarchical clustering to generate the heatmap. N = 3. (B) Venn diagrams showing the intersection between proteins identified in our study by OOPS and proteins annotated with RNA-binding GO terms (GO:0003723) downloaded from QuickGO (EMBL-EBI) for *Homo sapiens*. Duplicate UniProt accessions were removed in R prior to analysis. (C) Representative images of interphase RPE1 cells stained for PRC1 (in cyan) and DNA (Hoechst, in magenta) following treatment with RNase A. (D) Quantification of PRC1 nuclear fluorescence intensity from the staining shown in (C). Results are presented relative to the untreated control condition (RNase −). N = 3 (n_(no RNase)=_385; n_(with RNase)=_406). (E) Representative images of interphase RPE1 cells stained for SC35 (in cyan) and DNA (Hoechst, in magenta) following treatment with RNase A. SC35 staining is included as a negative control for the RNase treatment. (F) Quantification of SC35 nuclear fluorescence intensity from the staining shown in (E). Results are presented relative to the untreated control condition (RNase −). N = 3 (n_(noRNase)=_457; n_(with RNase)=_441). Error bars in (D) and (F) are shown as mean ± S.E.M. Scale bar, 5 μm. Nested t-test was used to determine statistical significance: ns, not significant; ***p < 0.001.

Interestingly, a subset of transient midbody proteins involved in cytokinesis, namely SHC Binding and Spindle Associated 1 (SHCBP1), Anillin (ANLN), and Protein Regulator of Cytokinesis 1 (PRC1), display RNA-binding activity predominantly in G1 and G1/S (Figure 6A). Consistent with our mass spectrometry data, western blots confirmed that although total protein levels of these factors rise and peak at G2/M (consistent with their roles in cytokinesis), their RNA-binding activity is maximal in early S and declines thereafter (Figure S8A, S8B). Despite well-characterised midbody functions during cytokinesis^63–66^ , their nuclear/interphase roles and the functional relevance of RNA binding remain poorly understood. We therefore focused on PRC1, which showed the strongest evidence of RNA-binding in the R-DeeP 3.0^10^ and RBR-ID^67^ datasets (Table S8). PRC1 nuclear staining in interphase cells was RNA-dependent (Figure 6C, 6D) as RNase treatment markedly reduced PRC1 signal. EDC4^68,69^, a P-body marker (Figure S8C, S8D), and SC35, a nuclear speckle marker (Figure 6E, 6F)^70,71^ , served as positive and negative controls for RNase treatment, respectively. These data identify PRC1 as a non-canonical RBP with peak RNA association at the G1-to-S transition and demonstrate that its interphase stability is RNA-dependent.

## Discussion

Understanding the regulatory principles of the cell cycle beyond well-studied protein networks is crucial for uncovering the mechanisms that control this fundamental process. In the present study, we have undertaken a systematic and comprehensive analysis of lncRNAs, transposable elements, alternative splicing, and RNA-binding proteins by integrating transcriptomic and proteomic profiling with the TEgment pipeline. To the best of our knowledge, our study provides the first RNA-centric atlas of cell cycle regulation in human normal diploid cells, using a palbociclib-based synchronisation strategy that minimises DNA damage.

First, our study showed that only 104 lncRNAs are cell cycle-regulated, far fewer than previously reported ^12–14,16^. This likely reflects differences in synchronisation agents, cellular models, the extent to which synchronisation drugs induce DNA damage and overall biological reproducibility across studies. These discrepancies highlight the need for caution when interpreting large, heterogeneous datasets and underscore the value of validation in diploid models and drug-independent synchronisation systems. Using PIP-FUCCI RPE1 cells, we confirmed that the poorly characterised lncRNA *ENSG00000230615* is enriched at the G2/M stage of the cell cycle, and that its depletion causes G1 arrest, indicating that loss of periodically expressed lncRNAs can impair cell cycle transitions. Because the depletion of chromatin-enriched lncRNA *ENSG00000230615* downregulated the adjacent *KLF17*, we propose that *ENSG00000230615* functions *in cis*. This aligns with reports indicating that *cis*-acting lncRNAs, which comprise a significant portion of lncRNAs with known functions, can influence the expression of nearby protein-coding genes^72^. Intriguingly, lncRNA *ENSG00000230615* is differentially expressed across multiple cancers, and pan-cancer analysis links high expression to worse survival, most notably in kidney clear cell carcinoma (KIRC) and adrenocortical carcinoma (ACC). Future studies should determine whether this syntenically conserved lncRNA mechanistically contributes to tumorigenesis and whether it may have prognostic value in these cancers.

A systematic analysis of TE expression (*Prokopov et al., 2026; manuscript co-submitted back-to-back*) using TEgment during cell cycle progression has not been reported previously. Although we observed differential expression of multiple TE classes across cell cycle stages, standalone AluS elements (SINE family^35,73^) are specifically enriched at the G1-to-S transition, suggesting a potential functional role in S-phase entry. The unique pattern of their periodic expression, combined with a prior report linking exogenously overexpressed AluS to S-phase progression^19^, indicates that AluS expression likely plays a functional role in the cell cycle. Given their S-phase–specific expression and the known ability of TE expression to activate oncogenes and promote tumorigenesis^74^, this raises the question of whether standalone AluS elements may rewire transcriptional programmes that drive uncontrolled proliferation and contribute to cancer.

Second, our data also establish that phase-specific groups of transcription (co)factors might be key controllers of the transcriptional landscape, driving orderly cell cycle progression. For example, the cell cycle expression of TEs suggested a common regulatory mechanism driven by evolutionarily recent KRAB-ZFP transcription factors. We identified two KRAB-ZFP associated with AluS regulation, namely ZNF334 and ZNF441, whose expression precedes Alu activation. This suggests that ZNF334 expression in G1 (T0) is associated with repression of AluS in that phase, as ZNF334 declines in G1/S (T5) while AluS levels increase. At the same time, we found that ZNF441 is specifically expressed in mid S (T8) while AluS and AluY, the latter known to be targeted by ZNF441^59^, are downregulated in G2/M (T11). This is consistent with the model in which KRAB-ZFP exhibit a family-wide tendency for synchronised cell-cycle expression that coincides with Alu silencing at corresponding cell-cycle stages^57^.

Third, we examined the regulation of alternative splicing across the cell cycle. The alternative splicing gene set defines a group not subject to periodic changes in steady-state transcript levels, confirming that splicing can be regulated independently of transcription during the cell cycle^18 6^. In contrast to the previous report^18^, we found that A5′SS, but not intron retention, was cell-cycle regulated, peaking at the G1-to-S transition, most likely due to differences in the computational design of the splicing tool^75^. By shifting exon boundaries, A5′SS can rapidly generate protein isoforms with altered domain composition^76,77^. For example, an A5′SS in *CEP63* produces a variant implicated in centriole duplication, a process required for accurate mitotic spindle assembly and faithful chromosome segregation, thereby illustrating how A5′SS may reshape the repertoire of centrosomal proteins during S phase. Among RBPs upregulated at G1-to-S transition, SRSF7 is a strong candidate regulator of the observed A5′SS enrichment. Reducing SRSF7 levels decreases the proportion of S-phase cells in cancer cells^62^, highlighting its importance during that phase. Although splicing regulation occurs at various levels^6,78,79^, our current analysis is transcriptomic. Future studies should include phosphoproteomics to understand how post-translational modifications of splicing factors contribute to cell cycle–dependent alternative splicing.

Our integrative analyses also showed that tRNA biogenesis may be regulated by the cell cycle, as we found TSEN54, a tRNA splicing endonuclease, enriched at the G1-to-S transition. Indeed, tRNA synthesis peaks during S phase^80^. Given the roles of specific tRNAs in proliferation and cell cycle arrest^81^, together with the well-documented alterations in tRNA expression in cancer^82^, we speculate that distinct tRNA pools and their extensive RNA modifications may be remodelled across the cell cycle to support stage-specific translational programmes that promote proliferation or contribute to tumorigenesis. Supporting this idea, we found that PUS10, a pseudouridine synthase involved in tRNA modification^83^, was enriched in the G1 phase of the cell cycle. However, we cannot exclude the possibility that PUS10 activity is also linked to cell cycle regulation of TEs, since loss of PUS10-mediated tRNA fragmentation has been associated with TE dysregulation^84^. Future work should profile the tRNA pools and their modification states across cell cycle stages and test their functional consequences for cell cycle progression and cancer phenotypes.

Finally, we also investigated the dynamics of RBPs across the cell cycle. Using OOPS ^26^, most RBPs that dynamically change RNA-binding activity are non-canonical, consistent with their multifunctional roles and the need to rapidly switch activities in response to cellular demand^9,85^. Intriguingly, several proteins with established roles in cytokinesis, namely Anillin, PRC1 and SHCBP1, are shown in this study to bind RNA during interphase. We further show that PRC1’s stability is RNA-dependent in interphase, where PRC1 displays nuclear staining. The moonlighting RNA-binding activities of cytokinesis proteins, their potential interphase functions, and whether these functions are linked to their cytokinetic roles remain largely unexplored. For PRC1, RNA binding could compete with microtubule binding, as shown for the microtubule-associated protein Tau^86^, or indicate roles in nuclear organisation and gene regulation, as proposed for nuclear Anillin^87^. Since PRC1 is deregulated in cancer and contributes to chromosomal instability^88^, it will be important to assess its nuclear functions, particularly those linked to RNA-binding activity, in oncogenic settings.

In summary, our study highlights the power of integrating transcriptomic and proteomic approaches to reveal how long non-coding RNAs, repetitive elements, alternative splicing, and RNA-binding proteins collectively orchestrate cell cycle progression. More broadly, our findings open new avenues for exploring RNA-mediated regulation of the cell cycle, with significant implications for biology and disease.

### Limitations of the study

This study used palbociclib-mediated synchronisation of RPE1 cells to profile RNA-based mechanisms across the G1, S and G2/M cell cycle stages. However, this approach does not distinguish between the G2 and M phases, limiting our ability to attribute changes specifically to mitosis. As a result, we cannot determine the specific contributions of lncRNAs, TEs, AS, and RBPs during mitotic progression. Additionally, although RPE1 cells are widely used as a model for studying cell cycle and mitosis, the generalisability of our findings may be restricted to this cell line. Examining the roles of lncRNAs, TEs, AS, and RBP dynamics in other normal diploid cell lines, and ideally in primary human cells, would help determine whether these regulatory patterns are cell line specific.

Although our bulk RNA-seq measures mature transcript levels and therefore reflects both synthesis and degradation, it does not directly assess nascent transcription, limiting inference about the transcriptional regulation of lncRNAs and TEs. Splicing analyses would also benefit from long-read sequencing to more accurately resolve isoforms and improve the detection of complex events. Given that our TF analysis relied on expression rather than chromatin accessibility, expanding the study to include stage-resolved changes in chromatin accessibility and TF motif analysis could reveal additional, accessibility-dependent TFs and illuminate regulatory dependencies not captured by expression alone. Finally, because both lncRNAs and TEs can be regulated by RNA modifications^89^, future work should interrogate the RNA-modifying enzymes and determine whether epitranscriptomic marks in their native RNA context oscillate across the cell cycle and influence cell cycle progression.

## Resource availability

### Lead contact

Requests for further information and reagents should be directed to and will be fulfilled by the lead contact, Lovorka Stojic.

### Material availability

This study did not generate new, unique reagents.

### Data and Code availability

All generated RNA-sequencing data are available in the GEO database (https://www.ncbi.nlm.nih.gov/geo/) under the accession number GSE322920. De novo transcriptome annotations used for TEgment analysis are accessible via Figshare DOI: 10.6084/m9.figshare.32064672. The mass spectrometry raw files and their associated MaxQuant output files are available in the PRIDE partner repository (http://www.ebi.ac.uk/pride/archive/) under accession number: PXD077622. This paper did not report any original code.

## Acknowledgements

We are grateful to all members of the Stojic lab for their comments and discussion of the manuscript. We would also like to thank Susana Godinho for the critical reading of the manuscript. We are thankful to the Cancer Research UK (CRUK) Barts Centre Microscopy Facility for their support with image acquisition, and to the CRUK Flow Cytometry, Proteomics, and Bioinformatics Core Facilities at the Barts Cancer Institute (BCI). We thank Jean Cook (University of North Carolina at Chapel Hill, USA) for PIP-FUCCI RPE1 cells. This work was supported by the Barts Charity (MGU0404, G-002906), Cancer Research UK Career Establishment Award (RCCFEL\100007), Royal Society Research Grant (RGS\R1\231139) and the Academy of Medical Science Springboard Award (SBF006\1026) to L.S. G.G. was supported by an AIRC Fellowship for Abroad. F.K.M. and M.D. are supported by a Cancer Research UK (CRUK) Programme Foundation Award (DRCPFA-Nov24/100005) to F.K.M. Ö.D., and D.P. were supported by Cancer Research UK Career Development Fellowship (RCCCDF-Nov21\100002). We acknowledge support from the CRUK City of London Centre of Excellence Award Core Funding to BCI (CTRQQR-2021\100004) for access to and support from core facilities.

## Author contributions

Conceptualization, L.S.; Investigation, G.G., A.O.C., C.B., P.B., M.D.C.R.R., M.D., and L.S.; Formal analysis, G.G., A.O.C., D.P., E.M., C.B., M.D.C.R.R., A.T., F.K.M. and L.S.; Data curation, G.G., D.P., E.M., and F.K.M.; Visualisation, G.G., A.O.C., D.P., E.M., and L.S.; Supervision, J. W., F.K.M., O.D., and L.S.; Writing – original draft, G.G. and L.S.; Writing – review & editing, all authors; Project administration, L.S.; Funding acquisition, L.S. All authors have read and approved the final version of the manuscript.

## STAR Methods

### Cell culture

hTERT-immortalised retinal pigment epithelial cells (hTERT-RPE1 / RPE1) were obtained from the American Type Culture Collection (ATCC; CRL-4000). PIP-Fluorescent Ubiquitination-based Cell Cycle Indicator RPE1 cells (PIP-FUCCI RPE1) were kindly provided by Jeanette Cook (USA). PIP-FUCCI RPE1 cells were cultured in Dulbecco’s Modified Eagle Medium (DMEM; Gibco, 41966-029), while RPE1 cells were cultured in DMEM/F12 (Sigma, D8437). Both media were supplemented with 10% fetal bovine serum (FBS; Gibco, A5256801) and 1% penicillin-streptomycin (Gibco,15140122). Cells were maintained at 37°C in a sterile, humidified incubator with 5% CO_2_. Both cell lines were regularly tested for mycoplasma contamination.

### Synchronisation of RPE1 cells with palbociclib

Palbociclib-mediated cell cycle synchronisation was carried out as reported^22^, with some modifications. hTERT-RPE1 were plated in 10- or 15-cm dishes at a density of 4×10^3^/cm^2^. The following day, cells were treated with 500 nM Palbociclib (Selleck, S1116) for 24 hours, then released by washing the dishes four times with warm medium for 3 minutes each. Cells were collected at the indicated time points and processed for downstream analysis.

### Sorting of PIP-FUCCI RPE1 cells

PIP-FUCCI RPE1 cells were trypsinised and pelleted by centrifugation at 400xg for 5 min RT. For sorting, cells were resuspended in DMEM supplemented with 3% FBS (Gibco, A5256801) and 2 mM EDTA (Thermo Fisher Scientific,15575020). A total of 300,000 cells in G1 (mVenus, 530/30 B-A), S-phase (mCherry, YG 610/20-A), and G2/M (mVenus and mCherry, 530/30 B-A and YG 610/20-A) were sorted using a BD FACSAria Fusion 4L (V-B-Y/G-R). Sorted cells were collected directly into TRIzol^TM^ LS (Sigma, T3934) for downstream RNA extraction.

### Subcellular RNA fractionation

RPE1 cells were grown in 15-cm dishes and processed to isolate RNA from cytoplasmic, nucleoplasmic, and chromatin fractions as described ^90^. The RNase-free Turbo DNase Kit (Thermo Fisher Scientific, AM1907) was used to remove DNA after isolating all RNA fractions. Total cellular RNA from the same RPE1 population was extracted directly using Trizol (Thermo Fisher Scientific, 15596026) and used as the normalisation reference for qPCR analyses. The use of total RNA for normalisation minimises potential localisation bias introduced by housekeeping genes. qPCR data were normalized using the following equation: % Input = 100 × 2^(Ct_total RNA − Ct_RNA fraction).

### RNA extraction, cDNA synthesis and qPCR

Total RNA was extracted from RPE1 WT cells using the RNeasy® Mini Kit (Qiagen,74106) and including a DNase I digestion step (Qiagen, 79254). For PIP-FUCCI RPE1 cells, RNA was extracted using TRIzol™ LS (Sigma, T3934) following the manufacturer’s protocol. cDNA was synthesised using the QuantiTect® Reverse Transcription Kit (Qiagen, 205313) according to the manufacturer’s instructions. qPCR was performed using PowerUp™ SYBR™ Green Master Mix (Applied Biosystems, A25742) on a QuantStudio™ 7 Flex Real-Time PCR System (Thermo Fisher Scientific). Thermal cycling conditions were as follows: 50°C for 2 min, 95°C for 2 min, followed by 40 cycles of 95°C for 1 s and 60°C for 30 s. *GAPDH* and *RPS18* were used as housekeeping genes in accordance with MIQE guidelines^91^. Primer sequences are listed in the Supplementary Information.

### Transfection of cells with LNA gapmers

RPE1 were seeded in 6-well plates and transfected at ∼50% confluency using Lipofectamine™ RNAiMAX (Thermo Fisher Scientific, 13778150) 24h after seeding. Antisense Locked Nucleic Acid (LNA) GapmeRs (QIAGEN) were used at final concentrations of 10 nM; these included a non-targeting control and two LNAs targeting lncRNA *ENSG00000230615* (Supplementary Information). Cells were harvested for RNA extraction or FACS analysis at 48h after transfection.

### Immunofluorescence

For the γH2AX staining, cells were seeded in 12-well plates on 12mm coverslips (VWR, ECN 631-1577), and synchronisation was performed as indicated above. At desired time points, coverslips were fixed with 4% paraformaldehyde diluted in PBS (Thermo Fisher Scientific, 28908) for 15 minutes at RT. Coverslips were then washed with PBS for 5 minutes, followed by permeabilisation with a solution of 0.5% Tween20 and 0.5% Triton X-100 (Acros Organics, 327371000) in PBS for 5 minutes at RT. Blocking was performed with 5% BSA for 30 minutes at RT, and primary antibodies were incubated overnight at 4°C (Supplementary Information). The next day, three 10-minute washes were performed with 0.1% Tween 20 in PBS (PBS-T), and then cells were incubated with secondary antibodies diluted in blocking buffer for 1h at 37°C. Three 10-minute washes were carried out again with PBS-T, followed by one wash with PBS before staining with 1ug/ml Hoescht diluted in PBS (Sigma, 33258) for 10 minutes at RT. Finally, coverslips were washed with PBS and water before mounting with ProLong™ Diamond antifade mountant (Thermo Fisher Scientific, P36961).

For the RNAse experiments, RPE1 cells were seeded in 8-well chambers (Ibidi, 80827) and mildly permeabilised with 0.3% Tween-20 for 10 min at RT. After permeabilisation, cells were quickly washed twice with PBS. RNase A (20mg/mL; Thermo Fisher Scientific, 12091021) was then added to designated wells, while PBS was used on the untreated controls. The samples were incubated for 20 min at 25°C. Cells were washed twice with PBS and then fixed with 4% paraformaldehyde as previously described. The following steps were the same as indicated above. At the end, instead of mounting, the wells of the 8-well chamber were washed 3 times with PBS for 10 minutes each, and the samples were stored in PBS, ready to be imaged.

### Cell cycle analysis by FACS

Cells were trypsinised, washed once with ice-cold PBS, fixed with 1mL of ice-cold 70% ethanol and then stored at 4°C. On the day of the analysis, fixed cells were centrifuged and washed twice with 2 mL of ice-cold PBS, resuspended in 500 µL of PBS, and incubated for 30 minutes at 37 °C with 50 µg/mL RNase A (TransGen, GE101-01). The cell suspension was filtered using 40μm cell strainers (FisherBrand, 22363547) and incubated for 30 minutes with 20 μg/mL propidium iodide (Sigma, P4170) on ice in the dark before proceeding with FACS analysis. Samples were run on LSR Fortessa (BD Biosciences), and 10,000 events were collected for each condition; data were analysed in FlowJo (v10.8.1).

### Western blot analysis

RPE1 cells were trypsinised, washed twice with ice-cold 1x PBS, and pelleted at 1,000 rpm for 3 minutes at 4 °C at the indicated time points. Pellets were lysed in NP-40 lysis buffer composed of 50 mM Tris-HCl pH 8 (15568025, Thermo Fisher Scientific), 125 mM NaCl (71386, Sigma-Aldrich), 1 % NP-40 (85124, Thermo Fisher Scientific), 2 mM EDTA (15575020, Thermo Fisher Scientific), 1 mM PMSF (93482, Sigma-Aldrich), cOmplete™ Mini EDTA-free protease inhibitor cocktail (11836170001, Roche) and phosphatase inhibitors (2 mM sodium fluoride, 201154, Sigma-Aldrich; and 1 mM sodium orthovanadate, S6508, Sigma-Aldrich). Samples were incubated on ice for 25 min and then centrifuged for 3 min at 12,000 x g at 4 °C. The supernatant was collected, and the protein concentration was estimated using the Bio-Rad Protein Assay (5000006, Bio-Rad). Proteins were denatured in 6x SDS buffer (0.375 M Tris pH 6.8, 12 % SDS, 60 % glycerol, 0.6 M DTT, 0.06 % bromophenol blue) at 95 °C for 5 min. Samples were loaded on Bolt® 4–12 % Bis-Tris Plus Gel (Thermo Fisher Scientific, NW04120BOX) and run in MOPS buffer (Thermo Fisher Scientific, B0001-02). Proteins were then transferred to a nitrocellulose membrane (1620115, Bio-Rad) using 1x Tris-Glycine transfer buffer with 20 % methanol and blocked with 5 % non-fat milk in TBS-T (50 mM Tris, 150 mM NaCl, 0.1 % Tween-20) for 1 hour at RT. The membranes were incubated with primary antibodies in 5 % milk in TBS-T at 4 °C overnight.

The membranes were washed 3 times the next day with TBS-T and incubated with horseradish peroxidase secondary antibodies (Agilent Dako, P0447 and P0448, dilution 1:2000). Immunobands were detected with SuperSignal™ West Pico PLUS Chemiluminescent Substrate (34580, Thermo Fisher Scientific), and the signal was developed using CL-XPosure films (34089, Thermo Fisher Scientific). Quantification of immunoblots normalised against appropriate loading controls was performed using ImageJ. The list of antibodies used for Western and uncropped scans of the immunoblots is provided in the Supplementary Information.

### Image acquisition and quantification

EDC4, PRC1, and SC35 staining were imaged using an Eclipse Ti-E inverted microscope (Nikon) equipped with a CSU-X1 spinning-disk confocal unit and a Zyla 4.2 sCMOS camera (Andor). The system included a motorised stage and Nikon Perfect Focus and was controlled using NIS-Elements software (Nikon). Imaging was performed with a CFI Plan Apochromat Lambda 60×/1.45 NA oil-immersion objective (Nikon), with optical sections acquired at 500nm. γH2AX stainings were acquired using an Olympus DeltaVision RT microscope (Applied Precision, LLC) equipped with a Coolsnap HQ camera. Image stacks were acquired in 0.5 μm steps, using Olympus x60 UPlanSApo oil immersion objective. Image deconvolution was performed with SoftWorx Explorer (Applied Precision, LLC).

#### Quantification of PRC1, SC35 and EDC4 signal

PRC1 and SC35 images were z-projected using the Sum Slices function in Fiji (v2.1.0)^92^. Nuclear fluorescence intensity was quantified using CellProfiler (v4.2.6)^93^. Nuclei were identified using the IdentifyPrimaryObjects module based on DAPI staining with minimum cross-entropy thresholding, followed by FilterObjects (AreaShape perimeter, eccentricity, and form factor) to remove poorly segmented nuclei. Fluorescence intensity was measured using MeasureObjectIntensity on the filtered nuclei.

#### Quantification of EDC4 foci

EDC4 images were analysed in Fiji (v2.1.0)^92^. Images were z-projected using Sum Slices and smoothed to reduce background noise. EDC4 foci per cell were quantified using the Find Maxima function with the prominence threshold set to 10,000.

#### Quantification of γH2AX foci

γH2AX images were analysed in Fiji (v2.1.0)^92^ . Images were z-projected using Max Intensity and smoothed to reduce background noise. Nuclei were segmented using the Analyse Particles function and γH2AX foci per nucleus were quantified using the Find Maxima function with a prominence threshold of 20.

### Pan-cancer analysis

Normalised RNA-seq gene expression values (FPKM-UQ), basic phenotype and survival data were retrieved from the GDC Pan-Cancer (PANCAN) dataset via the UCSC Xena Browser^28^. To remove lowly expressed genes, the dataset was filtered to retain genes with log_2_(FPKM-UQ + 1) > 1 in at least 10 cancer samples. Genes were mapped from Ensembl identifiers to gene symbols using the GENCODE human reference annotation^29^. Survival analysis was performed using the R packages survival (3.8-6) and survminer (0.5.2). For tumour-normal expression comparisons, normalised RNA-seq expression data (log_2_FPKM) were obtained from the TCGA-TARGET-GTEx dataset. Statistical significance was assessed using the Wilcoxon rank-sum test, with p-values corrected for multiple testing using the Benjamini-Hochberg method^30^. This paper does not report original codes. We have not generated any new software, algorithm, or unpublished custom code. We have provided references for the software used in this study.

### Orthogonal organic phase separation (OOPS) and total proteome processing

OOPS was performed as described previously^26^. Briefly, at the indicated time points, RPE1 cells were washed twice with ice-cold PBS and then irradiated on ice with 400 mJ/cm^2^ of UV-C (254 nm) using a Hoefer Scientific UV Crosslinker. Cells were lysed, scraped and collected in 2mL of Trizol LS (Sigma, T3934). For each mL of Trizol LS, 200 μl of chloroform was added, followed by vortexing. Phase separation was obtained by centrifuging 15 minutes at 12000xg at 4 °C. The interphase was isolated and washed further three times by resuspending in Trizol and repeating the phase separation by adding chloroform. The protein-RNA complexes were precipitated with 1mL of methanol and, after washing twice with methanol, the pellet was resuspended in 100 μl of RBP buffer (100 mM TEAB, 1 mM MgCl_2_, 1% SDS) and incubated 20 minutes at 95 °C. The RNA was digested with 2 μg RNase A/T1 (Thermo Fisher Scientific, 10775831) for 3h at 37 °C and then left to digest overnight at 37 °C upon the addition of a further 4 μg of RNase A/. The next day, phase separation was carried out as described above, the organic phase was collected, and proteins were precipitated overnight at -20 °C with 80% acetone. After washing twice with 80% acetone, the pellets were resuspended in 200 μl of 100 mM TrisHCl pH 8.8 8M Urea. A BCA assay (Thermo Fisher Scientific, 10678484) was performed to quantify protein concentration in the samples. As preparation for mass spectrometry, 20 μg of proteins was reduced with 10 mM DTT (Thermo Fisher Scientific, D1532) 30 minutes at room temperature and then alkylated with 55 mM iodoacetamide (G-Biosciences, 786-228) 30 minutes at room temperature (in the dark). Samples were loaded onto Vivacon 500 Hydrosart filters with a 30 kDa molecular weight cut-off (Sartorius, VN01H22) and centrifuged for 20 min at 14000xg. Samples were washed three times through the addition of 400 μl of UA buffer (8M Urea, 0.1 M Tris HCl, 8.5) to the filters, followed by centrifugation at 14000xg for 20 min. Samples were washed three times with 100 mM TEAB (Triethylammonium bicarbonate), and then protein digestion was performed overnight by incubating the membranes at 37 °C with 0.25 μg of trypsin (Sigma, T6567) diluted in 100 mM TEAB. TMTpro 18-plex labels (Thermo Fisher Scientific, A52047) were resuspended in 100% Acetonitrile, and peptide labelling was performed on the membrane with 5 μg of TMTpro labels per 1 μg of input for each sample filter. The labelling reaction was carried out for 1h at 25 °C with shaking at 600 rpm. The excess of TMT-Pro labels was quenched with 5% hydroxylamine and incubated for an additional 30 min at 25 °C. Peptides were eluted by centrifugation at 14000xg for 20 min, the filters were washed twice with 100 mM TEAB, and once with of 30% acetonitrile. All eluates (first elution + 3 washes) were collected, and at this stage, all samples were pooled. 100 μg of peptides was dried with a vacuum concentrator, resuspended in 300 μl of A* buffer (2% CAN, 0.1% TFA, 0.5% Acetic Acid), and 10μg processed for stage tipping^94^.

After stage tipping, the sample was dried with a vacuum concentrator, resuspended in 10 μl of A* buffer and 1.5 μg was injected for LC-MS/MS analysis.

For total proteome analysis, briefly, cells were trypsinised and pellets lysed in 200 μl of Tris-HCl pH 7.4 4% SDS buffer. Lysates were sonicated in a cold room with a tip sonicator: three 5-second pulses at an amplitude setting corresponding to 5 µm. Samples were cleared with Corning^®^Costar^®^Spin-X tubes (Merck, CLS8162-96EA) and then quantified with tryptophan assay. 25 μg of proteins was reduced and alkylated before processing with Vivacon® 500, Hydrosart filters, as already described above. Mass spectrometry sample preparation was performed as described for OOPS samples, except that, following peptide labelling, 100 µg of proteins were fractionated into seven fractions using the Pierce High pH reverse-phase fractionation kit (Thermo Fisher Scientific, 15362617) according to the manufacturer’s protocol. Fractions were dried using a vacuum concentrator and resuspended in A* buffer for LC-MS/MS analysis.

LC-MS/MS analysis was performed as described previously^95^ using a Q-Exactive plus Orbitrap mass spectrometer coupled with a nanoflow ultimate 3000 RSL nano HPLC platform (Thermo Fisher Scientific). An estimated 1 µg of each fraction or OOPS sample was injected into the mass spectrometer. Samples were resolved at a flow rate of 250 nL/min on an Easy-Spray 50 cm x 75 μm RSLC C18 column with 2 µm particle size (Thermo Fisher Scientific), using a 123 minutes gradient of 3% to 35% of buffer-B (0.1% formic acid in ACN) against buffer-A (0.1% formic acid in water), and the separated peptides were infused into the mass spectrometer by electrospray. The spray voltage was set at 1.95 kV and the capillary temperature was set to 255 _°_C. The mass spectrometer was operated in data-dependent positive mode, with 1 MS scan followed by 15 MS/MS scans (top 15 method). The scans were acquired in the mass analyser at 375-1500 m/z range, with a resolution of 70,000 for the MS and 35,000 for the MS/MS scans. Fragmented peaks were dynamically excluded for 30 seconds.

### Mass Spectrometry analysis

Raw MS files were searched against a FASTA file of the Homo Sapiens proteome, extracted from Uniprot (2023), using MaxQuant (version 2.2.0.0)^96^. Enzyme specificity was set to “Trypsin”, allowing up to two missed cleavages. Peptide and protein false discovery rates (FDR) were calculated using a reverse database search approach, and were both set to 1%. Reporter Ion settings for TMT 18plex were applied to the search, along with a minimum of 2 ratio counts per protein (Unique +Razor). All other settings were left as default. To remove batch effects from the TMT reporter intensity values, ComBat^97^ was used. All downstream data analyses, such as reverse and contaminants filtering, Log2 transformation, median normalisation, ANOVA analysis, Post Hoc test, and data visualisations, were performed in Perseus software^98^ (version 1.6.2.3).

### RNA-sequencing and splicing analysis

For RNA sequencing, four biological replicates of RPE1 cells were collected at each time point following cell cycle synchronisation with palbociclib. Libraries were prepared from 1 μg of purified RNA using RiboCop rRNA Depletion HMR V2 kit for ribosomal RNA depletion coupled to CORALL Total RNA-seq V2 kit (Lexogen, 183-184). Libraries were generated following the workflow for longer inserts (RLT chemistry), Lexogen UDI 12 nt (Lexogen, UDI12B_0001-0096) was used as an indexing system, indexed libraries were PCR-amplified and quantified with Qubit^TM^ dsDNA HS assay kit (Invitrogen, Q3285), finally, insert size was assessed with Tapestation (Agilent). Samples were pooled in equimolar ratios and sequenced to a depth of ≥ 40 million reads per sample with 150 bp paired end reads on NovaSeq 6000 (Novogene UK Company Limited, Cambridge).

The quality of raw reads was assessed with FastQC 0.11.9^99^ UMI codes were extracted using UMI-Tools 1.1.0^100^, and reads were trimmed with Trimgalore 0.6.5. Paired-end reads were aligned to the GRCh38.p13 human genome assembly using STAR 2.7.0 ^101^ with the two-pass mode. RSEM 1.3.1 ^102^ was used for transcript quantification, yielding gene counts which were imported into R using tximport 1.28.0 ^103^. DESeq2 1.40.2 ^104^ was used to perform differential expression analysis. Genes with at least ten counts in at least four samples were retained, resulting in 16015 genes for downstream analysis. A gene was considered phase-specific if i) it had the highest TPM expression in that phase compared with the other cell cycle phases, and ii) it was significantly differentially expressed in the adjacent phase, using a threshold of BH-padj < 0.05 and |log2FC| > 0.5. rMats 4.1.2^105^ pipeline was used to perform alternative splicing analysis using the STAR-aligned BAM files. The unpaired stats model was used to evaluate differences in alternative splicing events between sample groups. An alternative splicing event was considered phase-specific if i) it had the highest average inclusion level in that phase compared with the other cell cycle phases, and ii) |dPSI| ≥ 0.1 with FDR ≤ 0.05 compared with the adjacent cell cycle phase. Heatmaps and doughnut plots were generated using the R packages ComplexHeatmap(2.26.1) and ggplot2 (4.0.2).

### TEgment-based analysis of transposable element loci expression

TE loci were quantified and structurally classified using TEgment (*Prokopov et al., 2026; manuscript co-submitted back-to-back*). Reads were aligned to the GRCh38.p13 human genome with GENCODE v38 annotation using the TEgment workflow, which included read trimming (fastp 1.0.1^106^), *in silico* rRNA removal (SortMeRNA 4.3.7^107^), ^101^ 2.7.11b two-pass alignment, EASTR 1.1.2^108^ filtering of spuriously spliced alignments, reference-guided transcriptome assembly with StringTie 3.0.3^109^ in nascent-aware mode, and locus-level TE quantification with Telocal 1.1.1 ^110^. Differential TE expression analysis was performed in R using DESeq2 on locus-level TE count matrices. TE loci with a total count sum < 10 were excluded from downstream analysis. Pairwise differential expression was performed between adjacent cell-cycle phases (T0 versus T5, T5 versus T8 and T8 versus T11). A TE locus was considered phase-specific if i) it had the highest average variance-stabilised expression in that phase compared with the other cell-cycle phases and ii) it was significantly differentially expressed in the adjacent phase contrast using a threshold of BH-adjusted padj < 0.05 and |log2FC| > 0.5. Variance-stabilised counts were row-wise z-scored for visualisation. Heatmaps were generated using pheatmap 1.0.13 (https://github.com/raivokolde/pheatmap) after ordering loci by peak-expression phase and TE structural class, and temporal trend plots were generated using ggplot2.

### Filtering and analysis of phase-specific protein-coding genes

#### Identification phase-specific transcription factors (TFs) and cofactors (TcoFs)

Phase-specific TFs and TcoFs were identified by cross-referencing significant phase-specific protein-coding genes with the AnimalTFDB 4.0 database^43^. Only genes annotated as TFs or TcoFs in the human AnimalTFDB 4.0 catalogue were retained for downstream analysis. For each gene, log_2_TPM expression values were standardised across all samples by subtracting the gene mean and dividing by the standard deviation (z-score). The resulting phase-specific TF and TcoF lists were grouped by family and exported for visualisation. Heatmaps were generated for each cell cycle phase using *ggplot2* (version 4.0.2), each showing TF and TcoF expression across four time points (T0, T5, T8, T11) and four biological replicates per time point.

#### Analysis of TF activity by DoRothEA

Transcription factor activity across the cell cycle was inferred from the expression of significant phase-specific protein-coding genes using the DoRothEA regulon^48^ and the *decoupleR* R package (version 2.16.0). The DoRothEA human regulon was filtered to retain only high-confidence TF-target interactions (confidence levels A and B). TF activity scores were estimated using the multivariate linear model method implemented in *decoupleR*, requiring a minimum of 5 target genes per TF. TFs were considered significantly active if they displayed an absolute activity score greater than 2 and a nominal p-value below 0.05 in at least one sample. TFs with increased activity scores were selected for the final visualisation. Results were visualised as a heatmap using the *pheatmap* R package, with rows ordered by hierarchical clustering based on Euclidean distance and columns ordered by time point. Only increased activity scores are shown.

#### Identification of phase-specific splicing regulators and co-regulators

Phase-specific splicing regulators and coregulators were identified by cross-referencing significant phase-specific protein-coding genes against two databases. First, genes were annotated using the Gene Ontology (GO) database queried via the *biomaRt* R package (version 2.66.0) against the Ensembl database (GRCh38, release 115). Genes were retrieved using the following GO terms, grouped into three functional categories: core splicing factors (GO:0008380, GO:0000375, GO:0000377, GO:0000398, GO:0048024), splicing coactivators (GO:0071011, GO:0000245, GO:0000351, GO:0071004, GO:0071012, GO:0071013) and broader RBP-related terms (GO:0003723, GO:0003729, GO:0017069, GO:0030619, GO:0030620, GO:0016607, GO:0035145, GO:0000347, GO:0006397, GO:0034470, GO:0006396). Second, a curated list of human RBPs was obtained from the RBPDB database^60^ and mapped to Ensembl gene identifiers via *biomaRt*. When a gene was present in multiple categories, the most specific annotation was retained, prioritising core splicing factors over splicing coactivators, broader RBP terms and RBPDB-curated entries respectively. Duplicate entries were removed, retaining one record per unique Ensembl gene identifier. Finally, the gene selection was manually curated to retain only those with published evidence of a role in splicing regulation and/or RNA processing. For each phase-specific gene, log_2_TPM expression values were standardised across all samples by subtracting the gene mean and dividing by the standard deviation (z-score). Genes were grouped by cell cycle phase and ranked within each phase by mean z-score at their peak time point. Heatmaps were generated using *ggplot2* (version 4.0.2) to display expression across four time points (T0, T5, T8, T11) and four biological replicates per time point.

### Statistical analysis

All statistical analyses were performed using Prism 10 (v10.6.1) or R version 4.4.2. Data are presented as mean ± SEM from three biological replicates, unless otherwise stated. Statistical significance was defined as p<0.05. Details of statistical analysis and the number of replicates are provided in the figure and dataset legends. Superplots are used to show data distribution^111^.

