## Supplemental Information for "An RNA-centric atlas of human cell cycle regulation"

Supplementary information for this manuscript consists of:

- Supplementary Figures S1-S9
- List of LNA Gapmers, qPCR primers and antibodies used in the study
- Supplementary Tables Legends

A

| Study | Cell line | Syncronisation method | Biological Replicates | Type of study |
|---|---|---|---|---|
| *Ali et al. PMID 29491376* | HeLa | Thymidine + Hydroxyurea | One | S (nascent RNA-seq) |
| *Hao et al. PMID 33108271* | U2OS | Thymidine + Nocodazole | Two | G1,G1/S, S, G2, M (RNA-seq) |
| *Liu et al. PMID 28289232* | MCF7 | Hormone starvation, Thymidine, Nocodazole | Two/One | G0/G1,G1/S, M (GRO-, RNA-seq) |
| *Yildirim et al. PMID 32402276* | hTert-RPE1 | Mimosine | Two | S (nascent RNA-seq) |
| *Guiducci et al.* | hTert-RPE1 | Palbociclib | Four | G1, G1/S, S, G2/M (RNA-seq) |

F

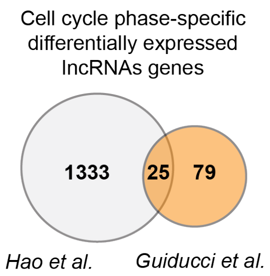


B
C
D
E

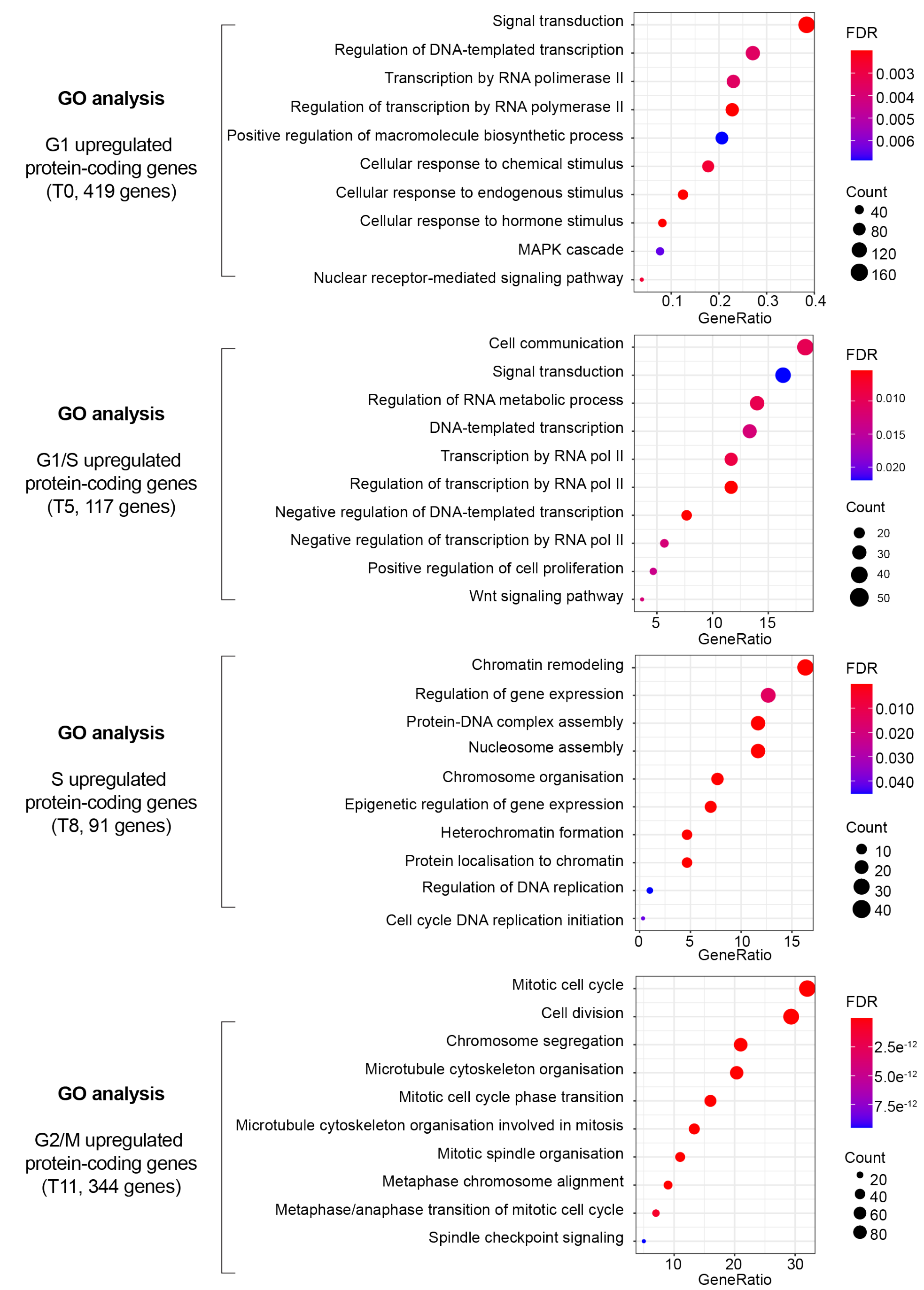

**Supplementary Figure S1 (Related to Figure 2). Cell cycle-regulated gene expression during palbociclib-mediated synchronisation of RPE1 cells**

(A) Summary of published studies investigating cell-cycle-regulated lncRNAs across different cellular models and synchronisation strategies. The present study is included.
(B-E) Gene Ontology (GO) analysis of protein-coding genes upregulated in the G1 (B), G1/S (C), S (D) and G2/M (E) stages of the cycle in RPE1 cells, identified by RNA-seq at the corresponding time points. The ten most relevant terms are shown.
(F) Venn diagrams showing overlap between cell cycle-regulated lncRNAs identified in this study (orange) and those reported by *Hao et al. 2020* [1] (grey).

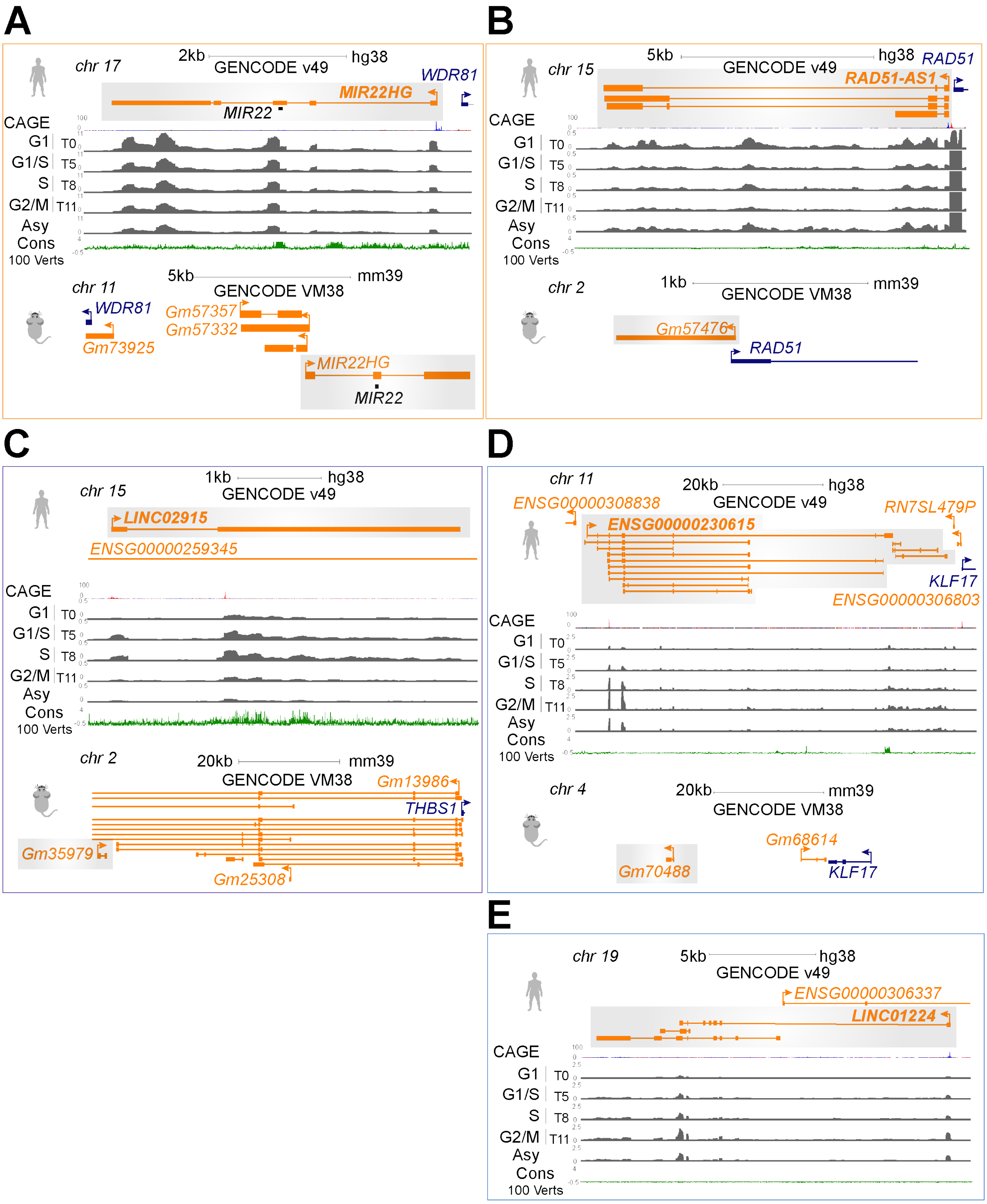


**Supplementary Figure S2 (Related to Figure 2). Genomic loci of selected cell cycle-regulated lncRNAs**

(A-E) Genomic loci of the G1 phase-enriched *MIR22HG* (A), *RAD51-AS1* (B), S phase-enriched *LINC02915* (C) and G2/M phase-enriched lncRNAs *ENSG00000230615* (D) and *LINC01224* (E). RPE1 RNA-seq coverage across the cell cycle is shown in grey, while FANTOM5 CAGE and conservation tracks were obtained from UCSC Genome Browser, with the corresponding orthologous mouse loci displayed. No mouse ortholog of *LINC01224* was detected.

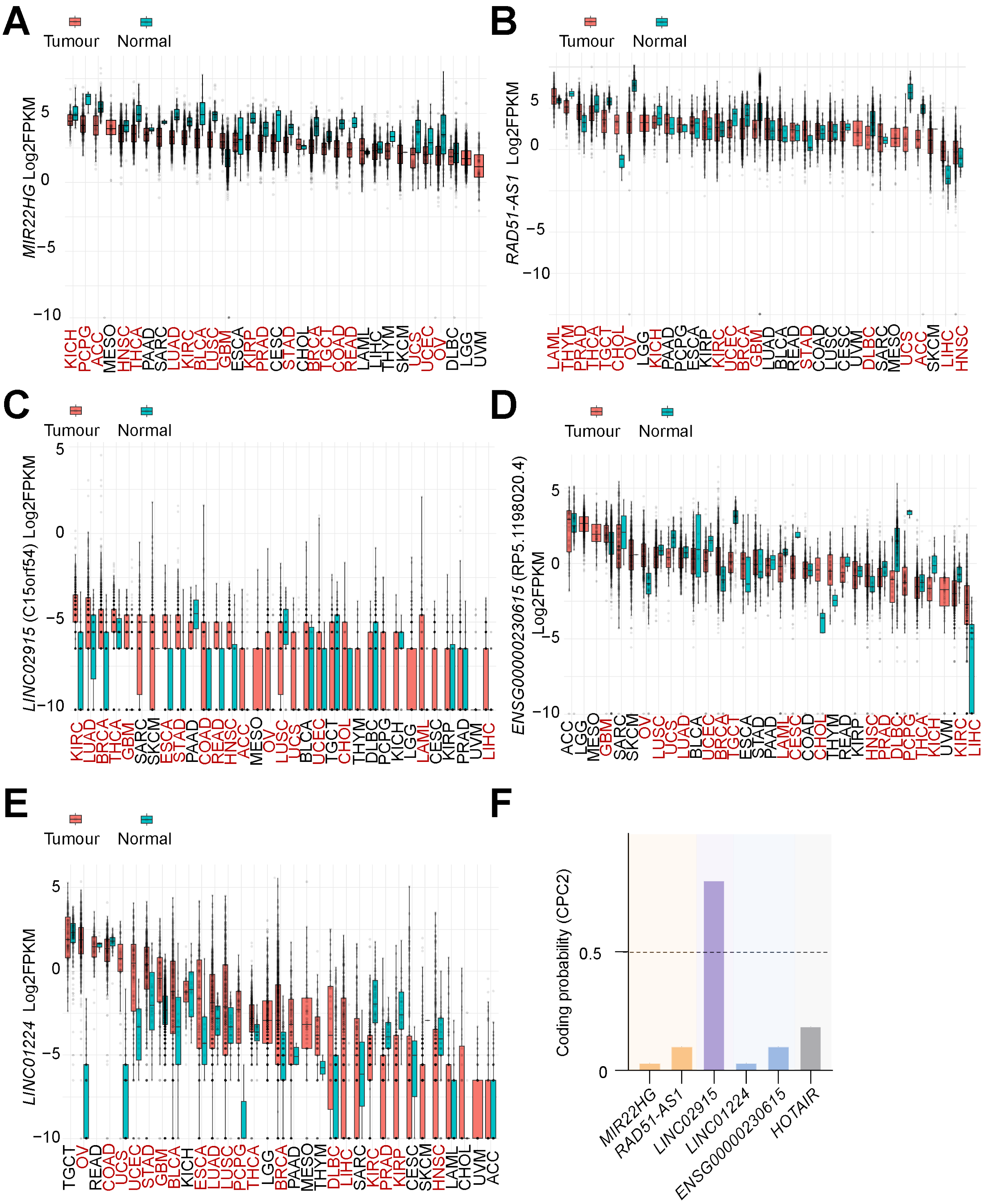


**Supplementary Figure S3 (Related to Figure 2). Coding potential and expression profiles of selected lncRNA candidates across normal and cancer tissues**

(A-E) Pan-cancer analysis of *MIR22HG* (A), *RAD51-AS1* (B), *LINC02915* (C), *ENSG00000230615* (D) *and LINC01224* (E) expression across the TCGA-TARGET-GTEx data. Boxplots of normalised log2(FPKM) gene expression is plotted for tumour vs normal tissues. Statistically significant tumour-normal comparisons are highlighted in red; corresponding p-values and sample numbers are provided in Supplementary Table S3. (ACC, Adrenocortical Cancer; BLCA, Bladder Urothelial Carcinoma; BRCA, Breast Invasive Carcinoma; CESC, Cervical & Endocervical Cancer; CHOL, Cholangiocarcinoma;

COAD, Colon Adenocarcinoma; DLBC, Diffuse Large B-cell Lymphoma; ESCA, Esophageal Carcinoma; GBM, Glioblastoma Multiforme; HNSC, Head and Neck Squamous Cell Carcinoma; KICH, Kidney Chromophobe; KIRC, Kidney Clear Cell Carcinoma; KIRP, Kidney Papillary Cell Carcinoma; LAML, Acute Myeloid Leukemia; LGG, Brain Lower Grade Glioma; LIHC, Liver Hepatocellular Carcinoma; LUAD, Lung Adenocarcinoma; LUSC, Lung Squamous Cell Carcinoma; MESO, Mesothelioma; OV, Ovarian Serous Cystadenocarcinoma; PAAD, Pancreatic Adenocarcinoma; PCPG, Pheochromocytoma & Paraganglioma; PRAD, Prostate Adenocarcinoma; READ, Rectum Adenocarcinoma; SARC, Sarcoma; SKCM, Skin Cutaneous Melanoma; STAD, Stomach Adenocarcinoma; TGCT, Testicular Germ Cell Tumour; THCA, Thyroid Carcinoma; THYM, Thymoma; UCEC, Uterine Corpus Endometrioid Carcinoma; UCS, Uterine Carcinosarcoma; UVM, Uveal melanoma). .

(F) Coding potential of selected cell cycle-regulated lncRNA candidates assessed by CPC2 [2] score. *HOTAIR* is included as an lncRNA with non-coding potential. The coding threshold is indicated by a dashed line.

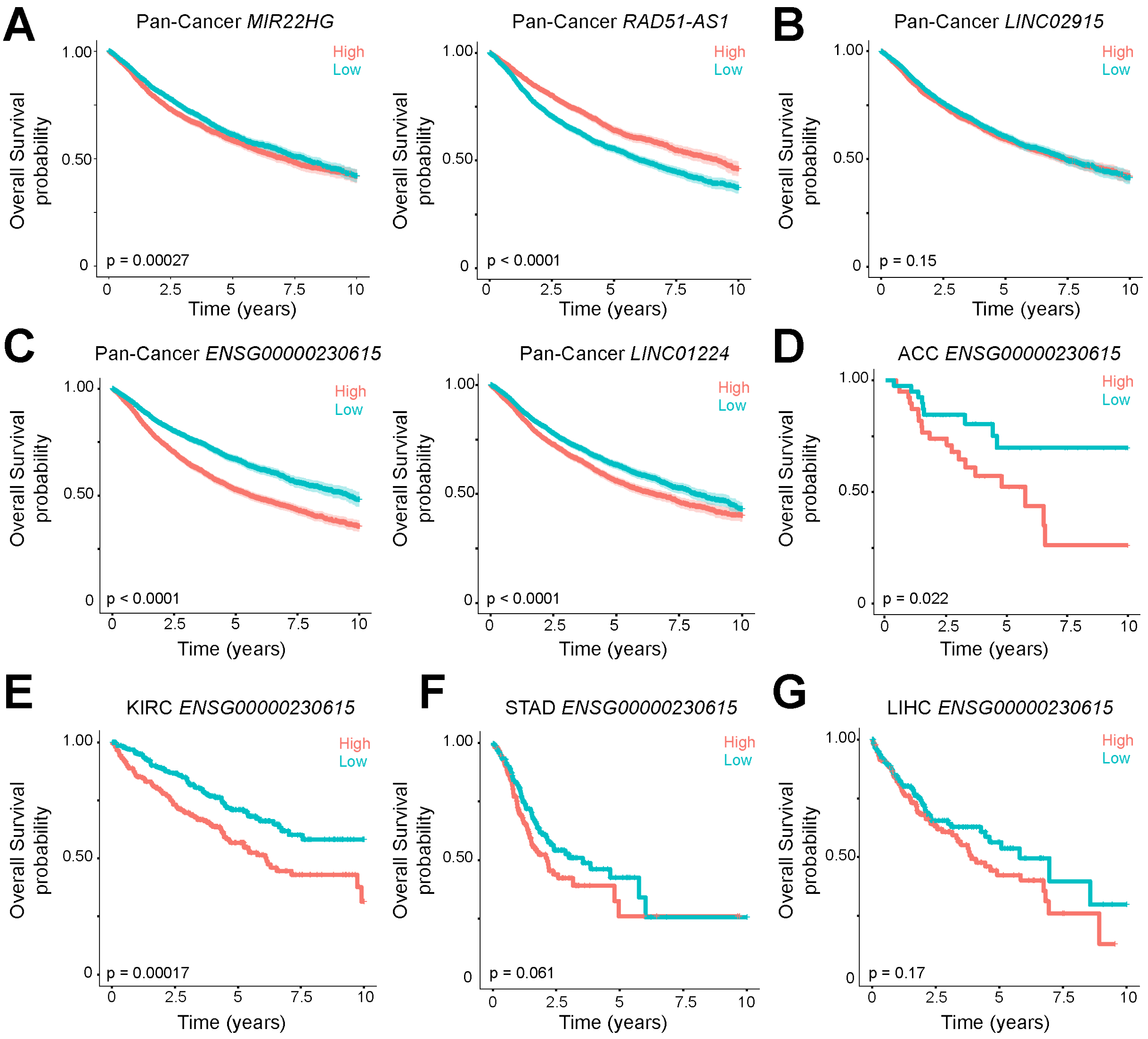


**Supplementary Figure S4 (Related to Figure 2). Survival curves of cell cycle-regulated lncRNAs from the TCGA-TARGET-GTEx dataset**

(A-C) Kaplan-Meier curves stratified by patients with high or low expression of *MIR22HG* (A), *RAD51-AS1* (A), *LINC02915* (B), *ENSG00000230615* (C) and *LINC01224* (C) from the TCGA-TARGET-GTEx dataset.

(D-G) Kaplan-Meier curves stratified by patients with high or low expression of *ENSG00000230615* in ACC (D), KIRC (E), STAD (F) and LIHC (G) from the TCGA-TARGET-GTEx dataset. (ACC, Adrenocortical Cancer; KIRC, Kidney Clear Cell Carcinoma; LIHC, Liver Hepatocellular Carcinoma; STAD, Stomach Adenocarcinoma).

High- and low-expression groups were defined by the median gene expression level. Log-rank p-values are indicated on the plots.

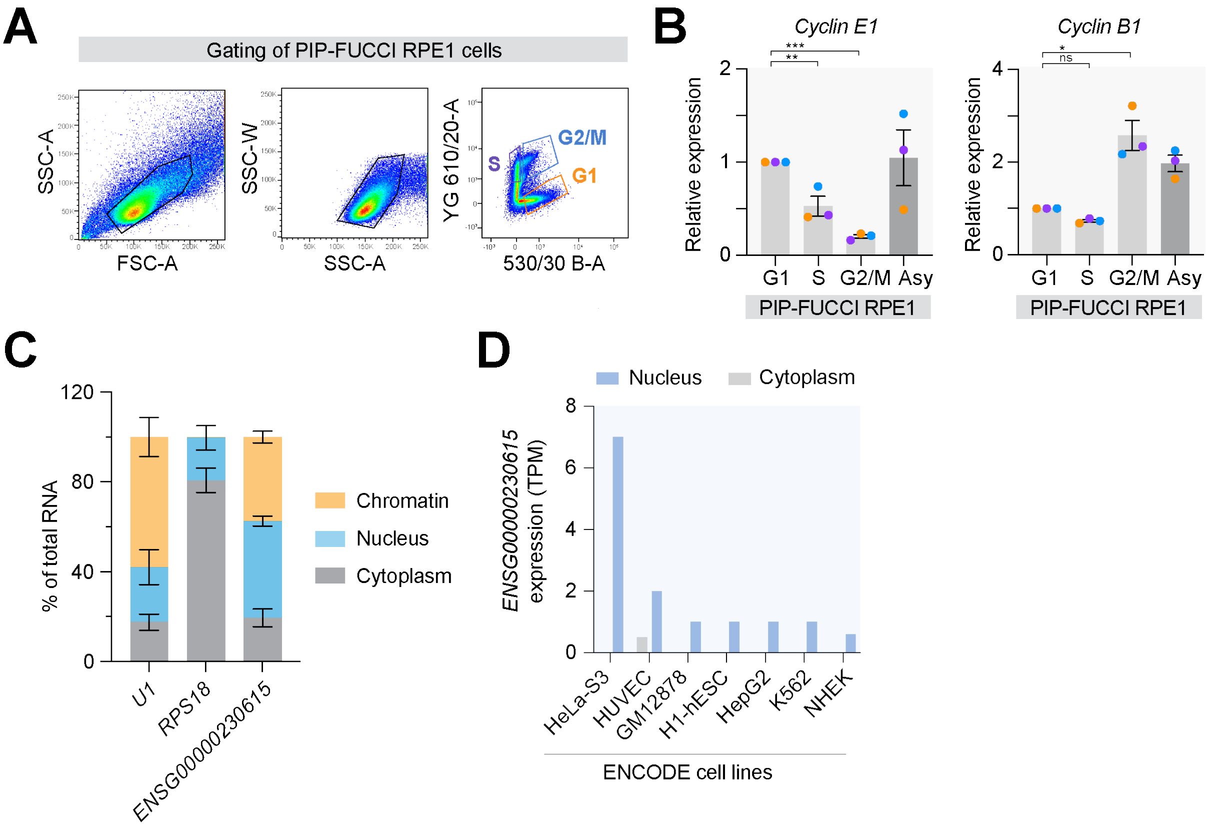


**Supplementary Figure S5 (Related to Figure 2). Gating of PIP-FUCCI RPE1 cells and characterisation of lncRNA *ENSG00000230615***

(A) Representative flow cytometry plots showing the gating strategy used to isolate PIP-FUCCI RPE1 cells at G1, S and G2/M stages of the cell cycle. The cells were first gated based on forward scatter area (FSC-A) versus side scatter area (SSC-A). Doublets were removed by gating side scatter width (SSC-W) vs SSC-A. PIP-FUCCI RPE1 cells were sorted based on cell cycle reporters YG 610/20-A versus 530/30 B-A.

(B) Expression of *cyclin E1* (left) and *cyclin B1* (right) in PIP-FUCCI RPE1 cells after sorting cells in G1, S and G2/M, as measured by qPCR. Results are presented relative to the G1 samples. N = 3.

(C) Expression of lncRNA *ENSG00000230615* in cytoplasmic, nucleoplasm and chromatin fractions of RPE1 cells. *RPS18* mRNA and *U1* small nuclear RNA were used as positive controls for the cytoplasm and chromatin fraction, respectively. N = 3.

(D) Expression of lncRNA *ENSG00000230615* in cytoplasmic and nuclear extracts of ENCODE cell lines, shown as transcripts per million (TPM) reads mapped. Data have been obtained from EMBL-EBI Expression Atlas (www.ebi.ac.uk).

Error bars in (B) and (C) are shown as mean ± S.E.M. One-way ANOVA with Dunnett's multiple comparisons test was used in panel (B) to determine statistical significance: ns, not significant; *$p < 0.05$; **$p < 0.01$; ***$p < 0.001$.

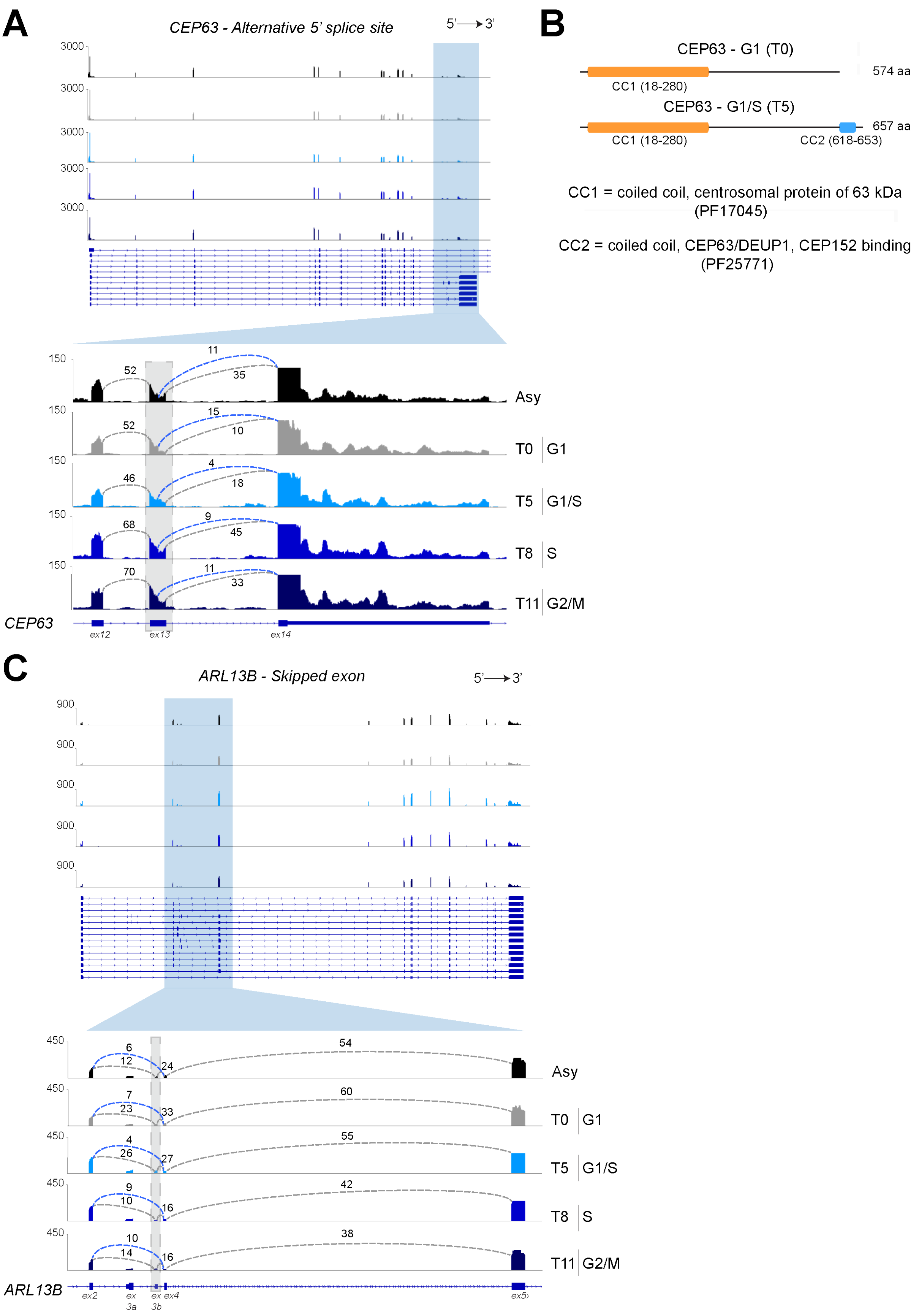
A
CEP63 - Alternative 5' splice site
5'→3'
3000
150
Asy
T0 G1
T5 G1/S
T8 S
T11 G2/M
CEP63
ex12
ex13
ex14
B
CEP63 - G1 (T0)
574 aa
CC1 (18-280)
CEP63 - G1/S (T5)
657 aa
CC1 (18-280)
CC2 (618-653)
CC1 = coiled coil, centrosomal protein of 63 kDa (PF17045)
CC2 = coiled coil, CEP63/DEUP1, CEP152 binding (PF25771)
C
ARL13B - Skipped exon
5'→3'
900
450
Asy
T0 G1
T5 G1/S
T8 S
T11 G2/M
ARL13B
ex2
ex 3a
ex 3b
ex4
ex5

**Supplementary Figure S6 (Related to Figure 4). Cell cycle-regulated alternative splicing of *CEP63* and *ARL13B***

(A) Sashimi plot showing an overview of the *CEP63* gene with a zoom-in of the alternative 5' splice site event in exon 13 observed at G1-to-S transition. The exon containing the alternative 5' splice site is highlighted with a grey rectangle. Tracks represent the averages of four biological replicates per condition: Asynchronous (black), T0 (grey), T5 and T8 (cyan and blue, respectively), and T11 (dark blue).
(B) Schematic of the two CEP63 protein variants detected during G1 and early S phase. The corresponding PFAM protein domains are indicated for each variant. Protein domain annotations were generated using InterProScan (https://www.ebi.ac.uk/interpro).
(C) Sashimi plot showing an overview of the *ARL13B* gene with a zoom-in of the skipped exon 3b in S-phase (T8). The skipped exon is highlighted with a grey rectangle. Tracks represent the averages of four biological replicates: Asynchronous (black), T0 (grey), T5 and T8 (cyan and blue, respectively), and T11 (dark blue).

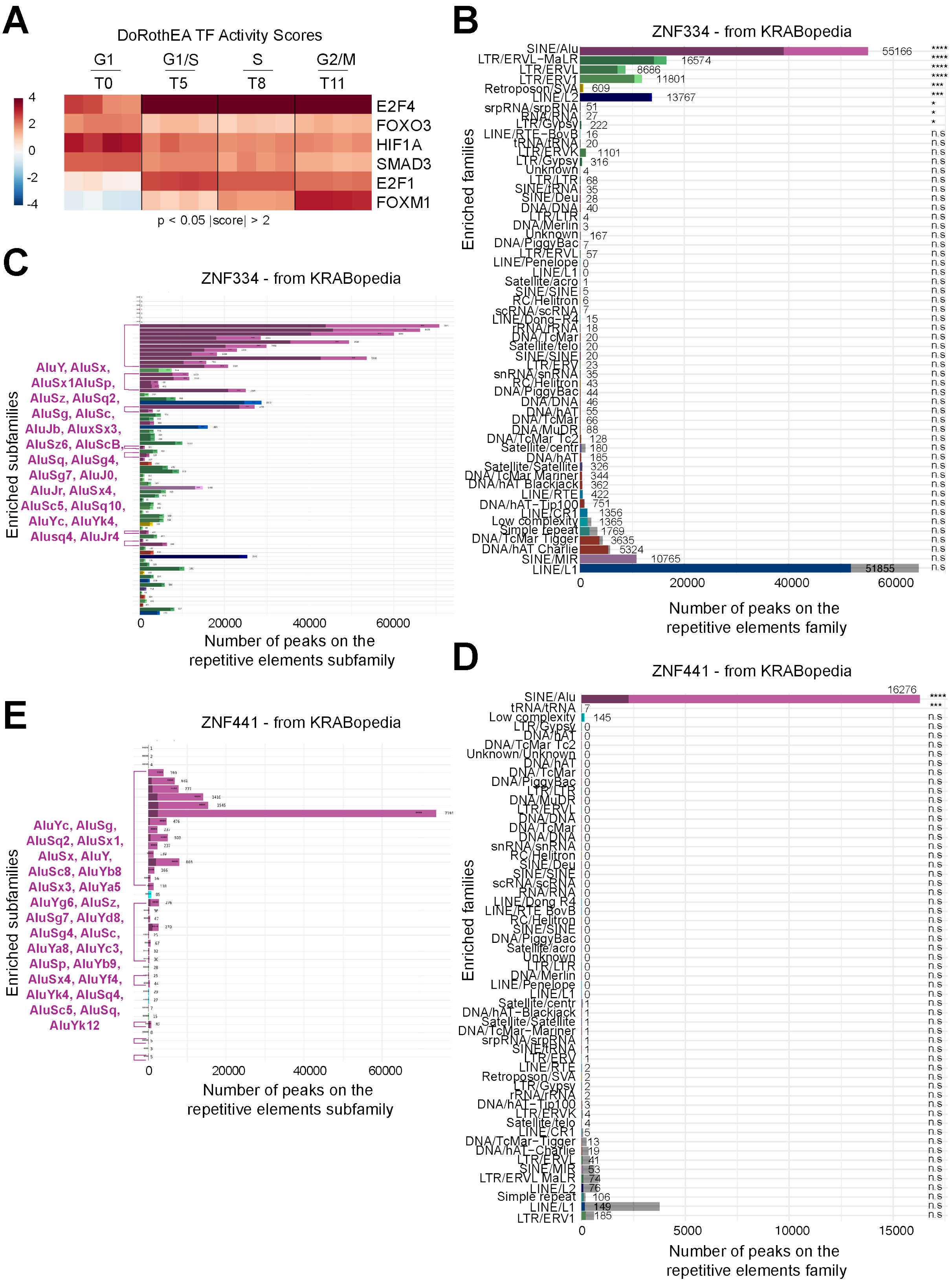


**Supplementary Figure S7 (Related to Figure 5). Integrated profiling of cell cycle transcription factor activity and KRAB-ZFP binding at transposable elements**

(A) Heatmap showing transcription factors (TFs) activity scores inferred from phase-specific differentially expressed protein-coding genes (Supplementary Table S1) using DoRothEA[3] (confidence levels A and B). Rows show TFs with significant activity changes (|score| > 2, p < 0.05 in at least one sample), while columns represent the four individual biological replicates ordered by timepoint (G1, early S, mid S, G2/M). Colour scale reflects normalised activity scores, only increased activity scores are shown (in red).

(B-E) ChIP-seq and/or ChIP-exo binding profiles of ZNF334 (B, C) and ZNF441 (D, E) retrieved from KRABopedia (krabopedia.org[4]). Bar charts show the number of peaks overlapping repetitive element families (B, D) and subfamilies (C, E), colour-coded by family. The darker portion of each bar represents the expected number of peaks following a random distribution. In the subfamily plots (C, E), Alu elements are highlighted in purple. Statistical significance of peaks enrichment at repetitive element families/subfamilies is indicated on the right (****FDR < 0.0001; ***FDR < 0.001; **FDR < 0.01; *FDR < 0.05; n.s. ≥ 0.05, not significant).

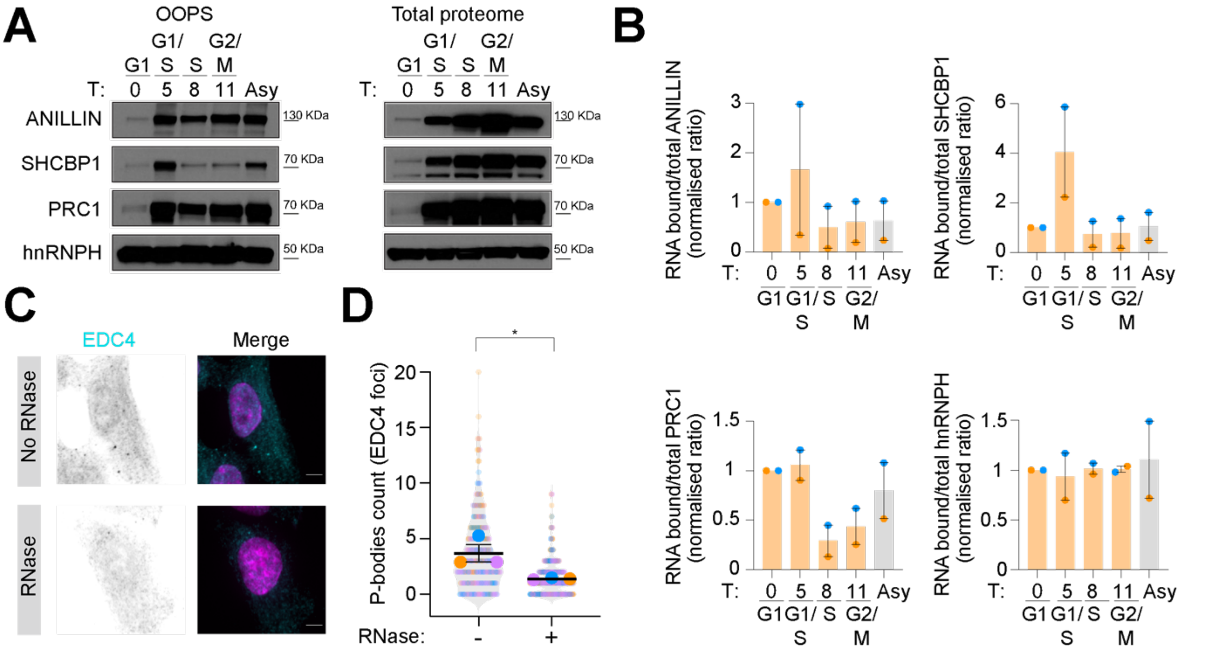


**Supplementary Figure S8 (Related to Figure 6). Validation of the RNA-binding activity of selected cytokinesis proteins in RPE1 cells**

(A) Western blot analysis of OOPS-derived and total protein fractions of selected cytokinesis proteins (i.e. ANILLIN, PRC1, SHCBP1) in RPE1 cells following palbociclib-mediated synchronisation. Time points corresponding to the G1, G1/S, S and G2/M phases are shown. hnRNPH was used as a negative control because its RNA-binding activity remains unchanged throughout the cell cycle.

(B) Densitometric quantification of normalised RNA-bound to total protein levels for cytokinesis proteins shown in (A). N = 2.

(C-D) Representative images of interphase RPE1 cells stained for EDC4 (cyan) and DNA (Hoechst, magenta) following treatment with RNase (C). Staining is shown alongside the corresponding quantification of EDC4 cytoplasmic foci, which are P-body markers (D). N = 3 ($n_{(RNAse-)}$= 312, $n_{(RNAse+)}$=331).

Error bars in (B) and (D) are shown as mean ± S.E.M. Nested t-test was used in panel (D) to determine statistical significance: ns, not significant; *$p < 0.05$.

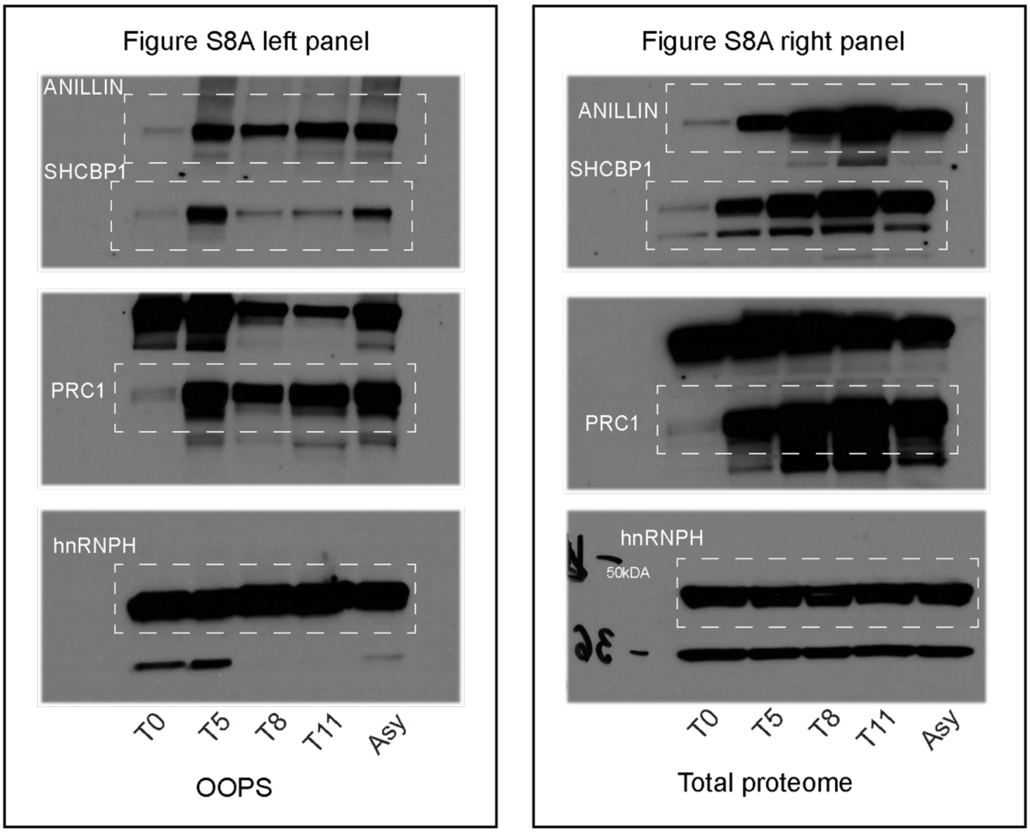


**Supplementary Figure S9. Uncropped pictures of the Western blot.** Regions in white boxes are shown in Supplementary Figure S8.

**List of antisense LNA gapmers (QIAGEN) used in this study**

| Target gene | Sequence 5' – 3' | Product number |
|---|---|---|
| LNA gapmer A<br>Neg control, QIAGEN | AACACGTCTATACGC | 300611-00 |
| LNA gapmer 1<br>*ENSG00000230615*<br>- Exon 2, QIAGEN | TCGGGAAAGAATTGAAG | 339511<br>LG00868997 |
| LNA gapmer 2<br>*ENSG00000230615*<br>- Exon 3, QIAGEN | ACGTGAGGACTGATAAC | 339511<br>LG00868999 |

**List of qPCR primers used in this study**

| Target gene | Forward primer (5' – 3') | Reverse primer (5' – 3') |
|---|---|---|
| *MIR22HG* | CGTTTTGGGCAAGTTGGAGA | CACATCTCGGAGGCAGAAAG |
| *RAD51-AS1* | GATCCTGCGCGAGTTTACAG | TCTTGGGAGTCGTGGTCTTC |
| *LINC02915* | ACACGTGCTGGATTCTCTCA | TCCTTTTAAACCCACACCGC |
| *LINC01224* | GTCCACAGGTTCTGCTAGCA | CAGGCCAAGGCAGAACAGTA |
| *ENSG00000230615* | CAGATGAGTCAGAGGTTTCTGC | CTCTTCTGCCGCATGTTTGG |
| *GAPDH* | CAACAGCCTCAAGATCATCAG | ATGGACTGTGGTCATGAGTC |
| *RPS18* | ATCCCTGAAAAGTTCCAGCA | CCCTCTTGGTGAGGTCAATG |
| *Cyclin B1* | TTTGCACTTCCTTCGGAGAGC | AAGGAGGAAAGTGCACCATGTC |
| *Cyclin E1* | TACACCAGCCACCTCCAGACAC | CCTCCACAGCTTCAAGCTTTTG |
| *KLF17* | CCACAGGCCAACAACAACAA | TCCTCTGAACAGTTGCCACG |

**List of antibodies used for immunofluorescence (IF) and Western Blot (WB).**

| Antibody | Supplier | IF Dilution | WB Dilution |
|---|---|---|---|
| EDC4 | 17737-1-AP, Proteintech | 1-200 | / |
| SC35 | S4045, Sigma-Aldrich | 1-2000 | / |
| PRC1 | 15617-1-AP, Proteintech | 1-400 | 1-1000 |
| Alexa Fluor® 488 donkey anti-rabbit | A21206, Thermo Fisher Scientific | 1-1000 | / |
| Alexa Fluor® 555 donkey anti-mouse | A21202, Thermo Fisher Scientific | 1-1000 | / |
| Cyclin B1 | 4138S, Cell Signaling | / | 1-1000 |
| Cyclin E1 | 20808S, Cell Signaling | / | 1-1000 |
| GAPDH | 2118, Cell Signaling | / | 1-2000 |
| hnRNPH | A300-511A, Bethyl | / | 1-1000 |
| ANILLIN | 48298, Cell Signaling | / | 1-1000 |
| SHCBP1 | 12672-1-AP, Proteintech | / | 1-1000 |

**Supplementary Table S1 (Related to Figure 2): List of significant phase-specific DEGs identified by RNA-seq and classified by phase.** Phase-specificity and statistical significance were calculated according to the following criteria: (i) highest TPM expression in one phase relative to other phases and (ii) differential expression with respect to adjacent phase (Benjamini–Hochberg -padj < 0.05 and |log2FC| > 0.5).

**Supplementary Table S2 (Related to Figure 2): GO analysis of significant phase-specific protein-coding genes identified by RNA-seq.**

**Supplementary Table S3 (Related to Supplementary Figure S3): Sample information and statistics for the comparison of selected lncRNAs expression in tumour vs normal samples** (TCGA-TARGET-GTEx).

**Supplementary Table S4 (Related to Figure 3): List of significant cell cycle-regulated TEs with corresponding TE classification and cluster assignment.** Cell cycle-regulated TE were defined as those differentially expressed between adjacent cell cycle phases with $|log_2FC| > 0.5$ and adjusted $p < 0.05$**.**

**Supplementary Table S5 (Related to Figure 4): List of significant phase-specific alternative splicing events identified by rMATS analysis.** Phase-specificity and statistical significance were calculated according to the following criteria: (i) highest average inclusion level (PSI) in one phase and (ii) significant differences relative to the adjacent cell-cycle phase (FDR < 0.05 and |dPSI| > 0.1).

**Supplementary Table S6 (Related to Figure 4): GO analysis of significant alternative spliced genes that are not differentially expressed.**

**Supplementary Table S7 (Related to Figure 6): OOPS-derived RNA-bound proteome intensities normalised to whole cell lysate-derived total proteome intensities across cell cycle stages.**

**Supplementary Table S8 (Related to Figure 6): RNA-binding evidence from RDeep3.0 (*Rajagopal et al*[5]*.*) and High-Resolution Mapping of RBPs (*He at al.*[6]) for proteins identified by OOPS in this study.**
